# Multimodal delineation of a layer of effector function among exhausted CD8 T cells in tumors

**DOI:** 10.1101/2023.09.26.559470

**Authors:** Arja Ray, Molly Bassette, Kenneth H. Hu, Lomax F. Pass, Tristan Courau, Bushra Samad, Alexis Combes, Vrinda Johri, Brittany Davidson, Katherine Wai, Patrick Ha, Grace Hernandez, Itzia Zaleta-Linares, Matthew F. Krummel

## Abstract

The anti-tumor function of CD8 T cells is limited through well-established pathways of T cell exhaustion (T_EX_). Strategies to capture emergent functional states amongst this dominant trajectory of dysfunction are necessary to find pathways to durable anti-tumor immunity. By leveraging transcriptional reporting (by the fluorescent protein TFP) of the T cell activation marker *Cd69,* related to upstream AP-1 transcription factors, we define a classifier for potent versus sub-optimal CD69+ activation states arising from T cell stimulation. In tumors, this delineation acts an additional functional readout along the T_EX_ differentiation trajectory, within and across T_EX_ subsets, marked by enhanced effector cytokine and granzyme B production. The more potent state remains differentially prominent in a T cell-mediated tumor clearance model, where they also show increased engagement in the microenvironment and are superior in tumor cell killing. Employing multimodal CITE-Seq in human head and neck tumors enables a similar strategy to identify Cd69RNA^hi^CD69+ cells that also have enhanced functional features in comparison to Cd69RNA^lo^CD69+ cells, again within and across intratumoral CD8 T cell subsets. Refining the contours of the T cell functional landscape in tumors in this way paves the way for the identification of rare exceptional effectors, with imminent relevance to cancer treatment.

**Graphical Abstract:** 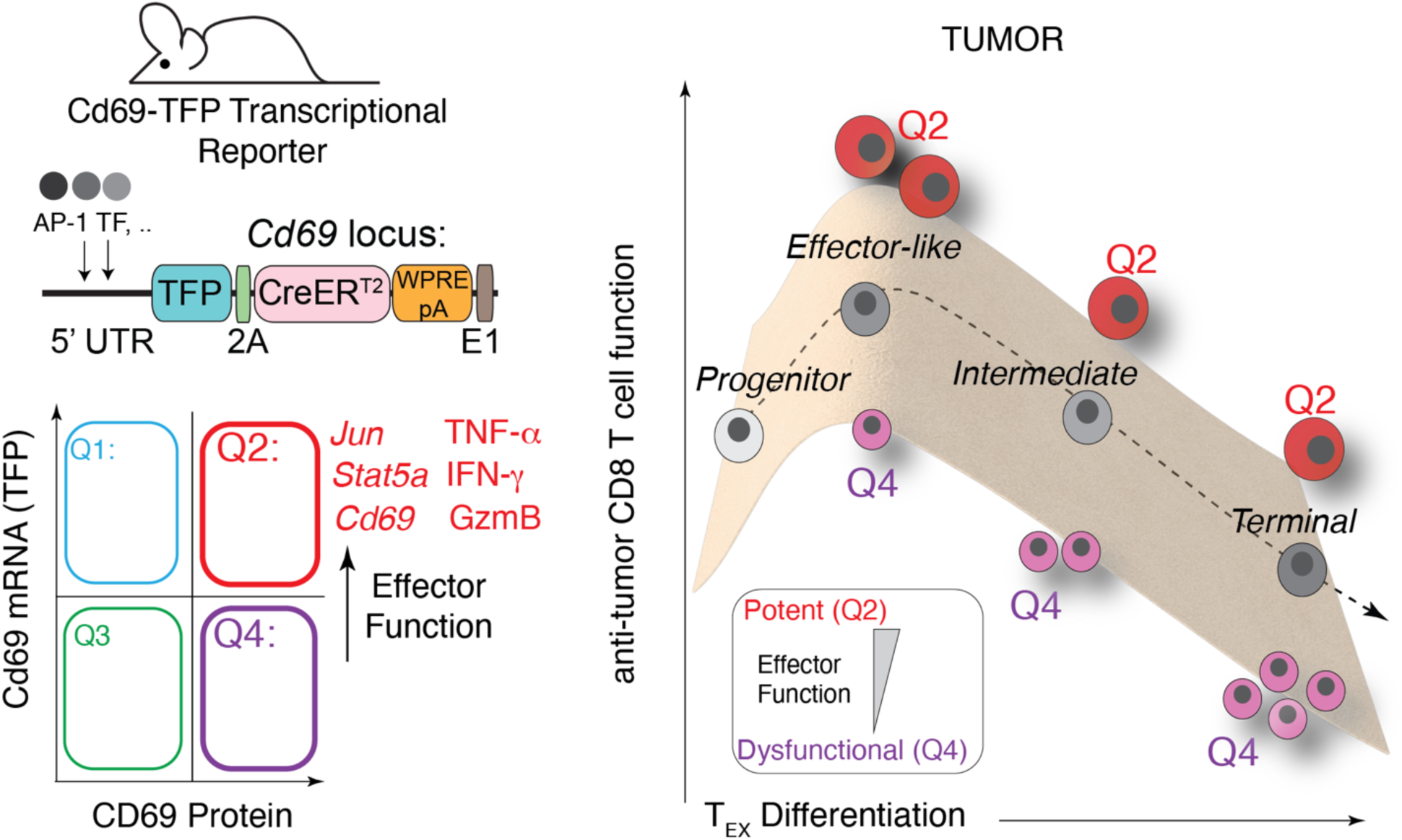

## Introduction

Within broadly immunosuppressive tumor microenvironments (TME), pockets of rare reactive immunity have been discovered, such as those containing conventional type 1 dendritic cells (cDC1s) that support CD8 T cells through antigen presentation(*1–4*). T cells, which integrate their encounters with antigens over their lifetime(*5–7*), require potent antigen stimulation for anti-tumor function. Yet chronic stimulation by persistent antigen in the TME conversely drives precursor CD8 T cells to dysfunctional or exhausted (T_EX_) states(*8*), notably driven by TOX expression(*9, 10*). This undesirable path to T cell exhaustion is increasingly well understood including its developmental stages(*11–13*), molecular markers(*10, 14–18*), transcriptional, epigenetic(*10, 15, 19–22*) and microenvironmental drivers(*23–26*). While such knowledge has been critical to shape our understanding T cell dysfunction in tumors, explicit strategies to delineate potent cytotoxic CD8 T cells within this exhausted milieu are not well-established.

Tissue-resident memory cells (T_RM_s) expressing CD69 and/or CD103 are one possible class(*27, 28*) that might diverge from the lineage of T_EX_ differentiation and define augmented effector function(*29*). While cells bearing these markers have been implicated in strong anti-tumor responses(*30, 31*), the degree to which cells defined by these markers are homogeneous or divergent from exhaustion are yet to be fully determined(*29, 32*). Efforts to identify potent effector functions in chronic stimulation conditions have also led to identification of their TCF1^hi^ progenitors, and the relevance of these to anti-tumor immunity is well-established, especially in the context of immune checkpoint blockade(*17, 33, 34*). However, such progenitor-like CD8 T cells can produce divergent trajectories of T_EX_ cells, including those biased towards effector-like states(*35*).

Substantial previous studies have mapped T_EX_ trajectories that lead to terminally exhausted T_EX_^Term^, notably identifying intermediates defined by KLR-expression (T_EX_^KLR^)(*12, 13*) which in turn have associated with better outcomes and cytotoxicity(*36*). Along the trajectory are also a population of Ly108-CD69-intermediate Tex(*37*), derived and distinct from the progenitor subsets that are Ly108+(*11*) and driven by TCF1 and MYB(*17*). These particular intermediate phenotypes likely also retain some plasticity to be directed away from terminal differentiation(*12, 36, 37*). Despite this well-delineated trajectory of differentiation, there is a diversity of cytotoxic potential among these T_EX_ subsets, and strategies to understand the potency of effector function within them are needed.

Evidence that effector function may not be fully dictated by the T_EX_ differentiation derives from studies of key transcription factors (TF), such as suppression of progenitor-associated factors TCF1(*19*) and MYB(*17*) expression, and over-expression of the AP-1 transcription factors JUN(*38*) as well as STAT5A(*37*) and BATF(*39*). Downregulation of early activation gene transcripts including *JUN* and *NR4A1* is a well-known response to stimulation in T and B cells(*40, 41*). AP-1 transcription factor over-expression on the other hand, counteracts the effects of TOX-mediated T cell dysfunction in the context of chronic viral infections and chimeric antigen receptor-expressing T cells(*37, 38*).

A key marker related to T cell function is the protein CD69 which has been used both to define T_RM_ identity and as a marker for early activation. Prior to our study, it was understood that T cells possess post-transcriptional regulatory mechanism for this ‘early-response’ gene which dissociates its transcription (associated with AP-1(*40, 41*) and therefore likely to decrease with repetitive stimulation) from its protein levels(*42, 43*). This dissociation of RNA and protein, which in a broader context is well-established in T and B cells(*44, 45*), offers an opportunity to study T cell states via comparisons of transcript levels related to history, separate from the CD69 protein levels indicating recent activation or long-term residency.

Here, we report the generation of a transcriptional reporter of CD69 RNA that together with paired analysis of CD69 protein, provides a novel window into the relationships between previous T cell activation history and their current capacity as cytotoxic and cytokine-producing effectors. The ability to mark such a dimension of effector states substantially refines the existing landscape of T cell dysfunction in tumors by providing a layer of gene expression and function, beyond that of the established differentiation trajectories.

## Results

While the cell surface expression of a protein like CD69 has historically been used to indicate both T cell stimulation and tissue retention(*46*), bioinformatic analysis of multiple datasets showed that the transcription of the *Cd69* gene was in fact inversely correlated with a history of chronic stimulation. For example, *Cd69* mRNA itself is higher in naïve vs. effector and early progenitor vs. terminally exhausted CD8 T cells **(Fig. S1a-c)**, is also higher in T cells in tumor-adjacent normal areas versus those within paired colorectal cancers (CRC) tumors (associated with T_EX_) (**Fig. S1d**). Further, expression of the transcription factors regulating *Cd69*(*47*) were also differentially lower in exhausted vs. naïve CD8 T cells (**Fig. S1e**). In contrast, CD69 protein expression, driven by TCR stimulation or other stimuli such as interferons(*48*), is often uncoupled from this transcriptional activity, as a result of strong 3’ UTR-mediated post-transcriptional regulation(*42, 43, 49*). We thus reasoned that tracking *Cd69* RNA alongside its protein might together provide a useful means to differentiate the potency of activation states in T cells.

### A novel genetic tool to report T cell stimulation history in vivo

To explore this idea, we generated mice in which DNA encoding the teal fluorescent protein (TFP), CreER^T2^ and a polyA site was inserted at the 5’ end of the *Cd69* locus (hereafter referred to as *Cd69*-TFP) **(Fig. 1a)**. In lymph nodes of unchallenged *Cd69*-TFP mice, the majority (∼80%, **Fig. 1b**) of CD8 T cells were TFP^hi^ without expressing surface CD69 protein, as measured by antibody staining (hereafter Q1 or TFP^hi^CD69-). We validated that the TFP levels we measured reflected *Cd69* RNA expression at steady state (**Fig. 1b, c**), and this is consistent with previous studies showing that resting T cells maintain relatively high levels of *Cd69* transcript that are translationally repressed until stimulation(*43, 50, 51*). In addition to this majority population, a small portion of CD8 T cells expressed CD69 protein on their cell surface alongside TFP (∼5%, Q2: TFP^hi^CD69+), possibly representing recently stimulated cells and a third population (∼15% Q3: TFP^lo^CD69-) was low for both TFP and CD69 (**Fig. 1b**). Both TFP and CD69 protein levels rose significantly in the context of robust stimulation - during the early and intermediate stages of thymic positive selection (**Fig. S2a-d**) (*52*) as well as during the first 3-16h of stimulation of isolated peripheral CD8+ T cells with anti-CD3/CD28 beads (**Fig. S2g**). CD69 protein positive cells in both of these settings, in which a majority of cells are expected to be antigen inexperienced, appeared dominantly in the transcript-high Q2 quadrant (**Fig. S2b, d**). We found that expression patterns for the CD69 protein were similar in reporter and WT mice in both cases (**Fig. S2f, i**), although surface CD69 levels by MFI were routinely about 50% lower (**Fig. S2e, h**). Difference in levels were stable when CD8 T cells were sorted by TFP expression and rested in IL-7 overnight suggesting they were not reliant on recent TCR stimuli (**Fig. 1d**).

**Fig. 1:**
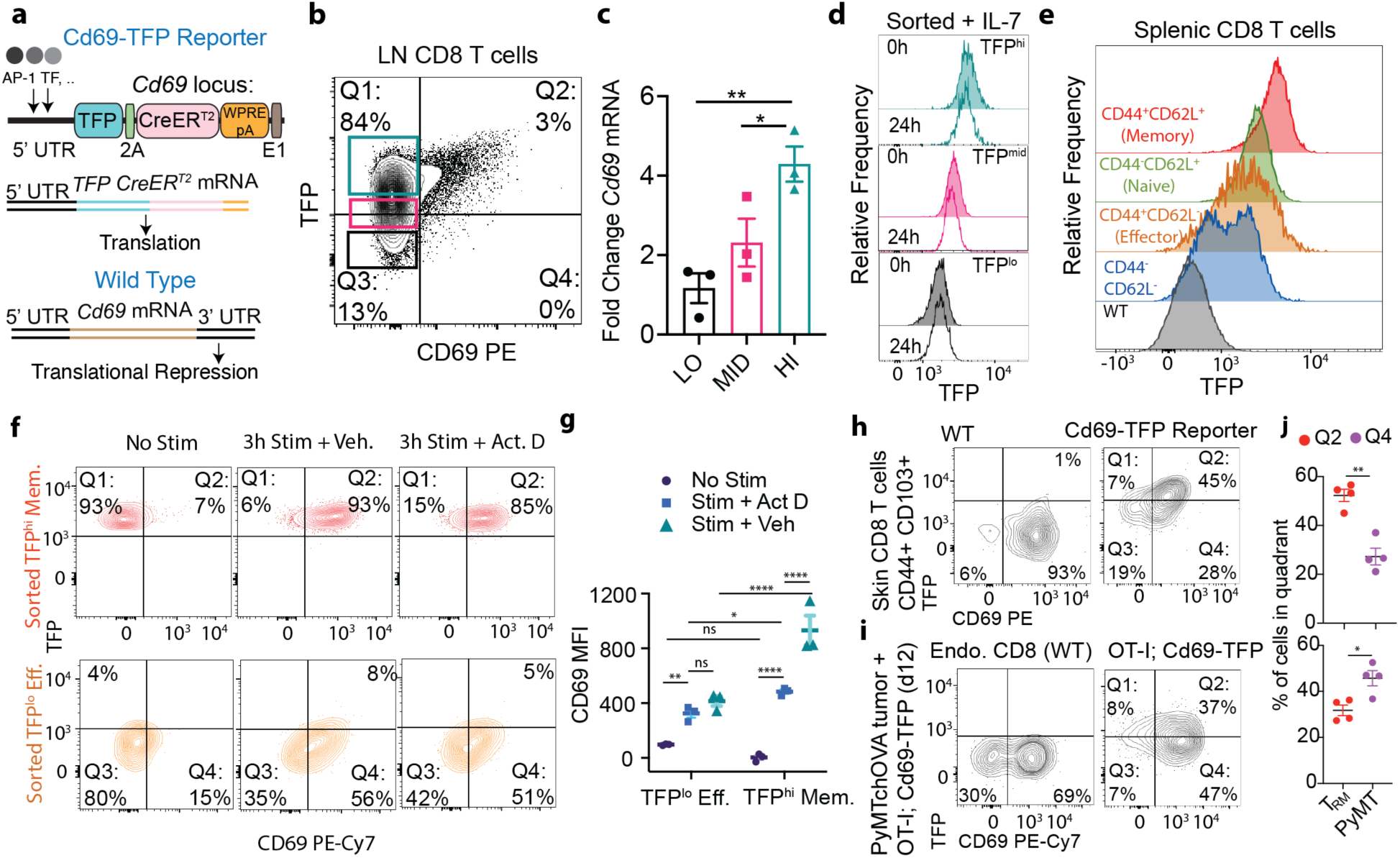
Distinct expression of Cd69 transcriptional reporter and CD69 protein in CD8 T cells. **(a)** Design of the TFP-2A-CreER^T2^-WPRE-pA reporter knocked into the 5’ UTR of the *Cd69* locus for transcriptional reporting; **(b)** TFP and surface CD69 expression in homeostatic lymph node (LN) CD8 T cells; color-coded boxes indicate sorted TFP ‘LO’, ‘MID’ and ‘HI’ cell populations, assayed for **(c)** Cd69 mRNA expression by qPCR; **(d)** histograms of TFP expression from sorted TFP^hi^ (top 20%), TFP^mid^ (middle 30%) and TFP^lo^ (bottom 20%) of splenic and lymph node-derived CD8 T cells at 0h and 24h post sort, resting in IL-7; **(e)** representative histograms of TFP expression in splenic CD8 T cells of different phenotypes (as indicated in the figure panel) from an unchallenged Cd69-TFP reporter mouse; **(f)** flow cytometry plots of TFP and CD69 in sorted TFP^lo^ Effector (CD44+CD62L-) and TFP^hi^ Memory (CD44+CD62L+) homeostatic CD8 T cells without stimulation (No Stim), 3h αCD3+αCD28 stimulation + DMSO (3h Stim + Vehicle) or 5μg/mL Actinomycin D (3h Stim + Act.D) and **(g)** CD69 MFI of the same across conditions; TFP and CD69 expression on **(h)** skin-resident CD44+CD103+ CD8 T cells in 6 month old unchallenged WT control and Cd69-TFP reporter mice; **(i)** Endogenous and adoptively transferred OT-I; Cd69-TFP reporter CD8 T cells in PyMTchOVA(*54*) tumors 12d post adoptive transfer and **(j)** corresponding quantification of the percentage of cells in Q2 (TFP^hi^CD69+) and Q4 (TFP^lo^CD69+) states in the same; Plots show mean +/− SEM; TFP negative gates derived from corresponding WT controls; null hypothesis testing by unpaired t test, adjusted for multiple comparisons, n=3 biological replicates representative of at least 2 independent experiments. *p<0.05, **p<0.01, ***p<0.001, ****p<0.0001, ns = no significance.

In unchallenged mice, TFP expression in CD8 T cells varied with differentiation state, consistent with Fig. S1a. Thus, CD44^lo^CD62L^hi^ naïve cells expressed higher TFP at baseline than CD44^hi^CD62L^lo^ effector CD8 T cells (**Fig. 1e)**. CD44^hi^CD62L^hi^ central memory T cells (memory) demonstrated higher levels still, which may be related to higher AP-1 transcription factor activity(*53*). CD44^lo^CD62L^lo^ cells expressed the lowest average level of TFP among the populations. When TFP^hi^ memory and TFP^lo^ effectors were sorted from spleens of unchallenged mice and stimulated with anti-CD3/anti-CD28 beads for 3h, TFP^hi^ cells generated more CD69 surface protein as compared to TFP^lo^ cells, an effect also observed in the presence of Actinomycin D treatment to block new transcription (**Fig. 1f, g**). Both groups maintained their pre-existing TFP status during this short stimulation, demonstrating how T cells occupying the Q4:TFP^lo^CD69+ cells may be generated from Q3:TFP^lo^CD69-cells (**Fig. 1f**). Finally, we found that the Q4: TFP^lo^CD69+ quadrant was modestly represented in skin-resident CD44+CD103+ CD8 T_RM_s from 6 mo. old mice (**Fig. 1h, j**) and dominant in ovalbumin-specific OT-I T cells in the spontaneous mammary tumors in genetically modified PyMTchOVA mice (*54*) 12d after adoptive transfer (**Fig. 1i, j**). The opposite was true for the Q2: TFP^hi^CD69+ quadrant (higher in T_RM_ than tumor) (**Fig. 1h-j**). Together with our informatic data, this supported the association between TFP levels and a history of encounters in CD8 T cells.

To further explore this possible relationship of TFP^lo^ cells with a history of repeated stimulation, we set up repetitive “chronic” stimulation cultures using purified CD8 T cells from *Cd69*-TFP mice. Cells were subjected to 3 cycles of 48h stimulation with 1:1 anti-CD3/anti-CD28 beads, followed by 72h rest under either hypoxia (1.5% O_2_) or ambient oxygen levels (normoxia), in the presence of low concentrations of IL-2 after the first cycle (**Fig. 2a, Fig. S3a**). The hypoxic condition has been shown previously to augment the acquisition of exhaustion-like features (*55*). *Cd69*-driven TFP levels at the end of each cycle were lower than the previous, an effect that culminated in about a 50% and 30% reduction under hypoxia and normoxia after 3 cycles, respectively (**Fig 2a-c, Fig. S3a, b**). Repeated stimulation concurrently upregulated exhaustion-associated markers such as PD1, CD38 and Tim-3, accentuated under hypoxia(*55*) (**Fig. 2f, Fig. S3d, e**). Addition of IL-2 without TCR stimulation (**Fig. S3f**) did not induce differentiation (**Fig. S3g**), decline in TFP expression (**Fig S3h**), nor acquisition of exhaustion-associated markers (**Fig. S3i**). While both *Cd69* mRNA (**Fig. 2d, Fig. S3c**) as well as the upstream AP-1 transcription factor *Jun* (**Fig. 2e**) decreased with repeated stimulation, a faster initial decay was observed when compared to the TFP reporter, likely due to longer half-life of TFP.

**Fig. 2:**
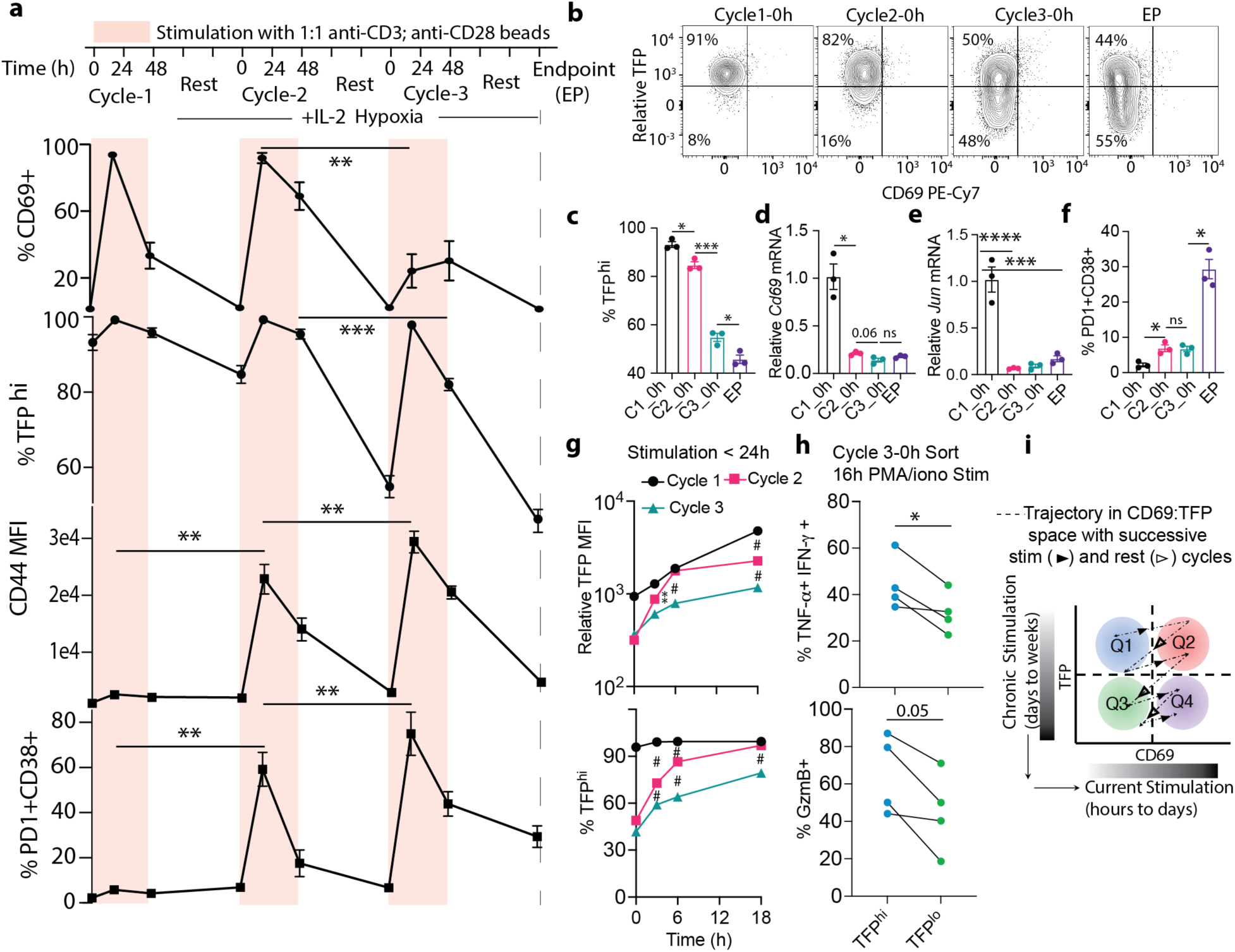
Repeated TCR stimulation drives down Cd69 transcription and the reporter TFP in CD8 T cells: **(a)** %CD69+, %TFP^hi^, CD44 MFI, %PD1^+^CD38^+^ of freshly isolated CD8 T cells through successive cycles of 48h stimulation and 72h resting in hypoxia + IL-2; **(b)** representative flow cytometry plots showing relative TFP (TFP normalized to WT control) and CD69 expression at the beginning of cycles 1, 2, 3 and endpoint (EP); **(c)** %TFP^hi^, **(d)** *Cd69* mRNA and **(e)** *Jun* mRNA by qPCR, **(f)** %PD1^+^CD38^+^ at the same time points; **(g)** TFP expression by MFI and %TFP^hi^ with 0, 3, 6 and 18h stimulation at Cycle1, 2 and 3h; **(h)** Cytokine and GzmB expression in sorted TFP^hi^ and TFP^lo^ cells from the ‘Cycle 3-0h’ condition, restimulated 16h with PMA/ionomycin; **(i)** schematic showing the trajectory of CD8 T cells within the TFP:CD69 states (quadrants) with successive stimulation and rest, providing a reading of history of stimulation. *p<0.05, **p<0.01, ***p<0.001, ****p<0.0001, ns = no significance, #p<0.001 in (g).

Analyzing the 24h time point in each cycle indicated a progressively lower magnitude of reporter expression by MFI, from Cycle 1 to Cycle 3 (**Fig. S3j**), which contrasted with effector differentiation and activation marker CD44, whose peak levels increased in every cycle (**Fig. 2a, Fig. S3a**). However, by the time we sampled 24h post stimulation, we noticed that %TFP^hi^ had reverted to ∼100% in each cycle (**Fig. 2a**). To test whether TFP levels after a shorter stimulation length is dependent on the history (cycle number), we further sampled stimulated cells at each cycle at 3, 6 and 18h time points (**Fig. 2g**). Indeed, both in terms of MFI as well as %TFP^hi^, we observed significantly and progressively lower levels of TFP expression at these intermediate time points (**Fig. 2g**). Therefore, to reach the same levels of TFP more stimulation is necessary in later cycles, demonstrating the inherent hysteresis and dampening in the system.

To test whether low TFP levels implicated different functionality, we sorted TFP^hi^ vs. TFP^lo^ cells from ‘Cycle3 0h’ and assayed cytokine and GzmB production after overnight (16h) PMA/ionomycin stimulation. This revealed that a higher proportion of TFP^hi^ cells and co-expressed TNF-α and IFN-γ and tended to be positive for GzmB compared to TFP^lo^ counterparts (**Fig. 2h**). A net model based on this set of experiments (**Fig. 2i**) proposes that in RNA (CD69:TFP) space, trajectories of CD8 T cell state are not retraced during subsequent activation events—rather, TFP levels decrease with repeated stimulation. Further, these in vitro experiments suggest that TFP^hi^CD69+ Q2 cells in this reporter system—displaying increased effector functions—are ones that are recently activated yet have not been subject to chronic and exhaustive stimulation.

### Delineation of chronic vs. potent activation states in tumors

To study this reporter in the context of *in vivo* tumor biology, we adoptively transferred *Cd69*-TFP reporter positive ovalbumin-specific CD8 T cells from CD45.1; OT-I TCR transgenic mice into WT mice bearing subcutaneously injected B78chOVA (OVA and mCherry expressed in B78, which is an amelanotic variant of B16F10 melanoma (*23*)) tumors (**Fig. 3a**). B78 is a an amelanotic B16 variant, which we have previously described as un-responsive to OT-I treatment and where adopted T cells are progressively rendered dysfunctional in the tumor(*23*). Recovered OT-Is from these tumors, were ∼90% PD1+ CD44+ (**Fig. S4a**) and were largely found in Q1 and Q2 of the reporter system on the first day they were detected (day 4), with endogenous intratumoral CD8 T cells used as the internal control to set the TFP gate (**Fig. 3b)**. While we found low levels of CD69+ among recently arrived T cells, consistent with the required loss of CD69 expression to emigrate from the dLN(*56*), by day 6-7 ∼70% of cells were TFP^hi^CD69^+^ (Q2: **Fig. 3b, d)**. By day 14, recovered cells were dominantly TFP^lo^CD69^-^ (Q3) and TFP^lo^CD69^+^ (Q4) (**Fig. 3b, d).** Consistent with our previous work using this(*23*) and other models(*54*), this shift to TFP^lo^ states was accompanied with increased differentiation towards chronic stimulation-driven exhaustion, here measured by PD1+CD38+(*15, 23*) among OT-Is (**Fig. S4b**), which also tracked with TOX^hi^TCF7^lo^ phenotypes (**Fig. S4c**). Similar trends in intratumoral OT-Is were observed in a spontaneous breast carcinoma tumor model (PyMTchOVA)(*54*) with adoptive transfer, albeit with a less prominent transient increase in Q2 (**Fig. S4d-f**).

**Fig. 3:**
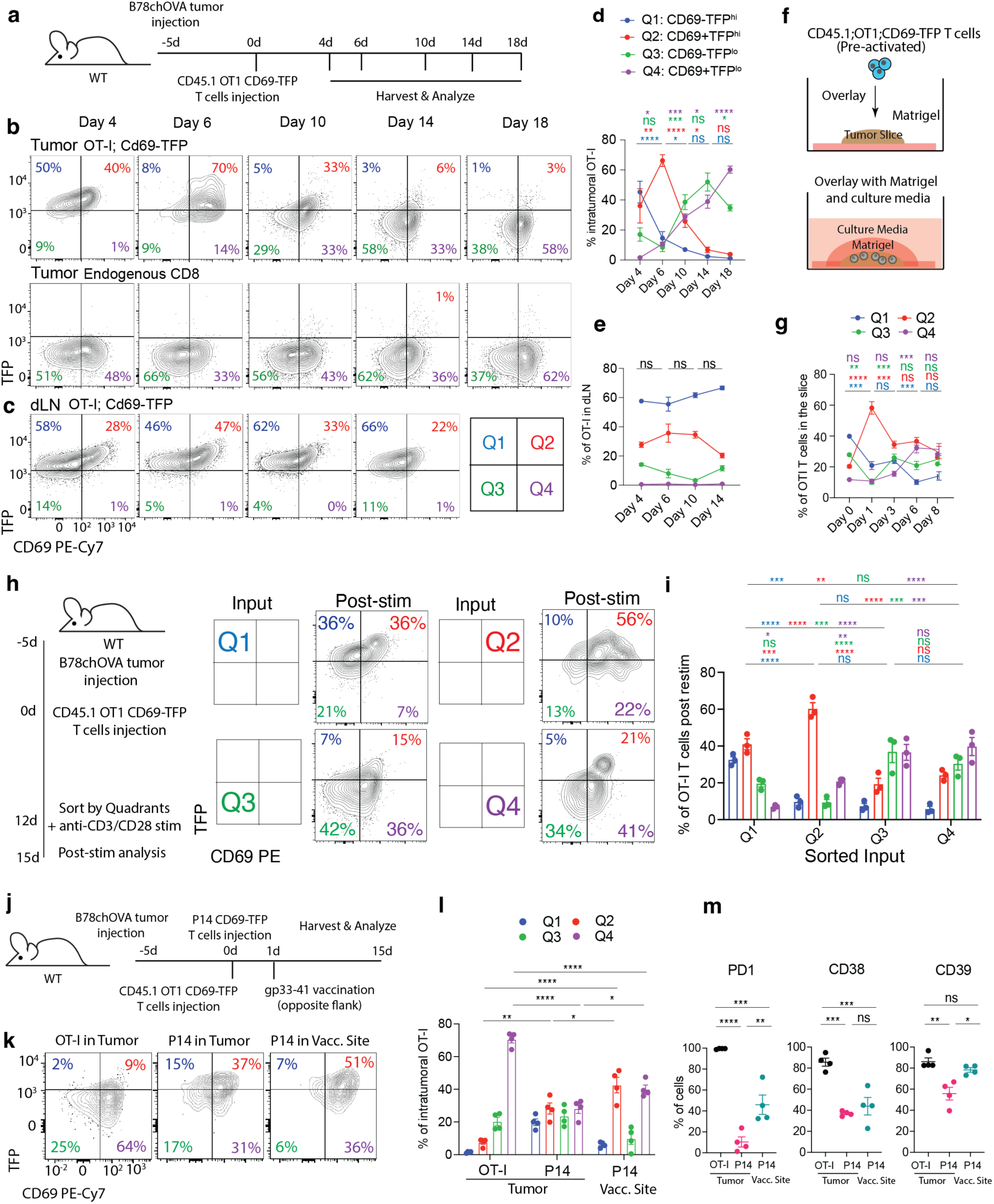
Chronic antigen exposure in tumor leads to decline in TFP expression in CD8 T cells. **(a)** Experimental schematic to track antigen-specific T cells in B78chOVA tumors over time; Flow cytometry plots showing TFP and CD69 expression of **(b)** adoptively transferred OT-I T cells and endogenous CD8 T cells from B78chOVA tumors and **(c)** OT-I T cells in corresponding tumor-draining lymph nodes (dLN) over time; corresponding CD69:TFP quadrant (Q1-Q4) distribution for the same OT-I T cells in the **(d)** tumor and **(e)** tdLN; **(f)** Schematic representation of long-term tumor slice culture setup; pre-activated: 48h stimulation with αCD3+αCD28 followed by 48h rest in IL-2; **(g)** CD69:TFP quadrant distribution of slice-infiltrating OT-I T cells over time; **(h)** representative flow cytometry plots and **(i)** quantification of resultant quadrant distribution of OT-I T cells sorted by their Cd69:TFP quadrants from B78chOVA tumors (Q2-Q4) or dLN (Q1) 12d post adoptive transfer, and restimulated with anti-CD3/anti-CD28 for 3d; **(j)** Experimental schematic of tumor injection and contralateral vaccination with distinct antigen specificities (OT-I and P14; P14 cells are 20-25% less TFP+ than OT-Is at baseline before injection); **(k)** Flow cytometry plots showing TFP and CD69 profiles of OT-I, P14 T cells in the OVA+ tumor and P14 T cells at the gp33-41 vaccination (vacc.) site; **(l)** CD69:TFP quadrant distribution of the same and **(m)** PD1, CD38 and CD39 expression among the same three groups. Representative data from 2-3 independent experiments, 3-4 mice or 5-6 slices/timepoint/experiment, pre-slice-overlay samples in duplicate, plots show mean +/− SEM. ****p<0.0001, ***p<0.001, **p<0.01, *p<0.05 in one-way ANOVA for each quadrant with pairwise multiple comparisons as shown using the post-hoc Tukey test in j, k, and 2-way ANOVA with post-hoc Tukey test between pair-wise timepoints (d-g) and 2-way ANOVA with false discovery rate correction for comparisons among inputs (i).

An alternative hypothesis to attributing these changes to differentiation is one in which cells migrating from the lymph node were progressively decreasing in TFP levels and replacing those in the TME. When the draining lymph node (dLN) was analyzed, in contrast to the tumor, we found the vast majority of cells remaining TFP^hi^ throughout the experiment. (**Fig. 3c, e**) suggesting that the observed changes more likely relied on features of the tumor microenvironment. As another way to address the competing hypothesis more directly, we modified a tumor slice overlay protocol(*57*) to create an experiment in which a selection of T cells encounter the TME at the same time with no further migration being possible (**Fig. 3f**). In this setting, the progression of phenotypes through CD69:TFP quadrants (**Fig. 3g, Fig. S5a)** and the acquisition of exhaustion markers (**Fig. S5b-d**) recapitulated what we had seen in vivo, suggesting that encounter with the tumor microenvironment (TME) was sufficient to drive these changes. Slice cultures also allowed analysis of proliferation in the slice-infiltrating OT-I T cells over time using violet proliferation dye (VPD) (**Fig. S5e**). VPD dilution accompanied the general decrease in TFP expression with each division, exemplified at day 3 **(Fig. S5f**). Q4 cells at d8 had undergone more division (**Fig. S5g**) and were significantly more PD1^+^CD38^+^ than those from Q2 (**Fig. S5h**), suggesting that they were further along the T_EX_ differentiation trajectory. In fact, analyzing the level of VPD within cells in the four CD69:TFP quadrants at each time point reinforced the trajectory of transition from TFP^hi^ Q1 and Q2 states to TFP^lo^ Q3 and Q4 states (**Fig. S5i**).

To further assess the conversion of T cells among these CD69:TFP states in the context of tumors, we sorted OT-Is d12 post transfer by their CD69:TFP phenotypes from B78chOVA tumors (Q2-Q4) and the dLN (Q1), restimulated these cells in vitro with anti-CD3/CD28 beads, and tracked the resultant quadrant transitions after 3d (**Fig. 3h, i**). Q1 cells transitioned to Q1 and Q2 and partially to Q3, while Q2 converted to Q2 and Q4, with small percentages in Q1 and Q3 (**Fig. 3h, i**). Sorted Q3 and Q4 cells both remained predominantly (∼70-80%) within the Q3 and Q4 states – suggesting a propensity to inter-convert between these states (**Fig. 3h, i**). Of note, both Q3 and Q4 cells retained some ability to convert to TFP^hi^ Q2 upon strong stimulation, although this propensity was significantly lower than those of Q1 and Q2 cells (**Fig. 3h, i**). Together, these data reveal a dominant trajectory of conversion from TFP^hi^ to TFP^lo^ states in tumors, consistent with T cells undergoing repeated TCR stimulation.

To further determine how this progression from TFP^hi^ to TFP^lo^ states is related to antigen stimulation and the corresponding microenvironment, we isolated CD8 T cells with a non-tumoral specificity (LCMV-specific P14; *Cd69*TFP) and assessed their state both within a tumor that did not express their antigen or from within a vaccination site, as compared to OT-I T cells from an OVA-expressing tumor (**Fig 3j**). When the P14; *Cd69*TFP cells were co-injected with CD45.1; OT-I; *Cd69*TFP T cells into B78chOVA tumor-bearing mice, that then received a priming gp33-41 peptide vaccination distal to the tumor (**Fig. 3j**), we found that they expressed higher TFP levels in the tumor than OT-Is (**Fig. 3k, l**). P14 T cells at the contralateral vaccination site also remained substantially TFP^hi^ with a 5x increase in the frequency of cells Q2 (**Fig. 3k, l**) than tumor OT-Is. These differences in CD69:TFP quadrant distributions aligned with the expression of surface markers PD1, CD38 and CD39, associated with exhaustion (**Fig. 3m**). Overall, these surface proteins were highest in the intratumoral OT-Is, intermediate in the P14s at the vaccination site and comparatively low in the non-specific intratumoral P14s (**Fig. 3m**). Hence, exposure to the TME alone did not lead to loss of a Q2 state, and presentation of antigen at a different site also did not lead to decreases in TFP level. Overall, these data support that antigen-specific stimulation of T cells specifically in a TME can drive a CD69+TFP^lo^ (Q4) phenotype.

### TFP^hi^CD69+ Q2 marks a state of relative functional potency in T_EX_ cells

We next sought to establish whether Q4 and Q2 cells were also functionally different in tumors and determine how this related to T_EX_ cell differentiation in tumors, best highlighted in single-cell sequencing(*12, 58, 59*). We thus pooled OTI; *Cd69* TFP T cells from eight B78chOVA tumors 12d after adoptive transfer and sorted for CD69:TFP quadrants, barcoded each population separately and performed single cell RNA Sequencing (scSeq) (**Fig. 4a**). From 13,352 cells, nine computationally derived clusters were identified (**Fig. 4b**), driven largely by the same canonical markers of previous works and with similar predicted differentiation trajectories (**Fig. S6a-c**). Strong associations were observed between the gene expression patterns corresponding to Tex^Prog^, Tex^E.Eff^, Tex^Term^, Tex^ISG^ and Tex^KLR^ subsets in our dataset and those that previously established this nomenclature(*12*) (**Fig. S6e**). This is consistent with the use of the same nomenclature in this context, although some differences exist such as in the T_EX_^Int.^ and T_EX_^Mem^ subsets – and may be attributable to variability between tumor and LCMV-derived subsets in those other works.

**Fig. 4:**
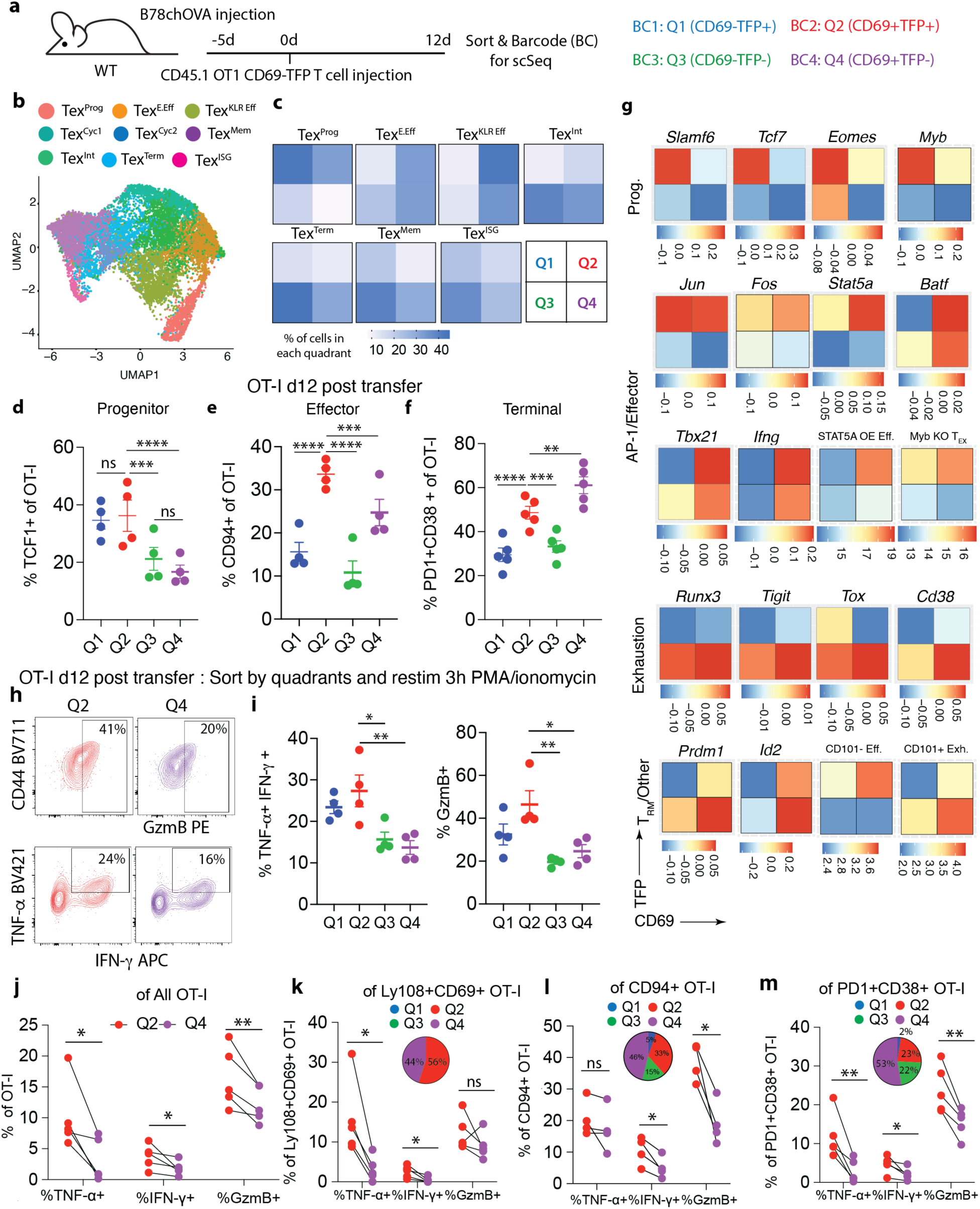
TFP^hi^CD69+ (Q2) marks a potent functional state of antigen-specific CD8 T cells within tumors: **(a)** Experimental schematic for single cell transcriptomic profiling of intratumoral OT-I T cells sorted by CD69:TFP quadrants; **(b)** UMAP representation of the scSeq data color-coded by computationally-derived clusters; **(c)** Heatmap showing percentage of cells that belong to the four CD69:TFP quadrants Q1-Q4 within each cluster; distribution of **(d)** percentage of cells bearing progenitor (TCF1+), **(e)** effector (KLRD1, or CD94+) and **(f)** terminal exhaustion (PD1+CD38+) markers within each CD69:TFP quadrant; **(g)** Heatmaps showing distribution of transcripts (canonical progenitor, AP-1/Effector, Exhaustion and tissue-resident memory related genes) and transcriptomic signatures (corresponding to Stat5a overexpressing effectors(*37*), Myb KO(*17*), CD101-Tim3+ and CD101+Tim3+(*13*) exhausted CD8 T cells) in the four CD69:TFP quadrants; **(h)** flow cytometry plots showing representative GzmB and cytokine expression in Q2 vs. Q4 intratumoral OT-Is from B78chOVA tumors 12d post adoptive transfer which were sorted and then restimulated for 3h in PMA/ionomycin and **(i)** corresponding bar graph quantification of the same; **(j)** cytokine and GzmB expression in intratumoral OT-I T cells at baseline (d12 post adoptive transfer) in the Q2 vs. Q4 activation state; cytokine and GzmB expression of Q2 vs. Q4 OT-I T cells within **(k)** Ly108+CD69+(*11*), **(l)** CD94+ and **(m)** PD1+CD38+ intratumoral OT-I subsets with inset pie chart showing the percentage of cells in each quadrant) within the corresponding subsets. Bar plots show mean +/− SEM; null hypothesis testing by ratio paired t test (i-m), RM ANOVA followed by post-hoc paired t-tests corrected for multiple comparisons (d-f). *p<0.05, **p<0.01, ns = no significance.

Barcoding of sorted cells by CD69:TFP states then allowed us to map quadrant distributions of these post-exhaustion T cell subsets (T_EX_)(*12*), (**Fig. 4b, c, Fig. S6d**). In agreement with the evolution of CD69:TFP quadrant distributions in Fig. 3, T_EX_^Prog^ cells were barcoded as coming from to Q1 or Q2 states and more terminally differentiated subsets of T_EX_^Int^, T_EX_^Term^ were dominantly derived from Q3 and Q4 barcoded cells (**Fig. 4c**). T_EX_^KLR Eff^, previously the most likely candidates to have high effector function, were heterogenous, expressing barcodes indicative of either Q2 or Q4.

Using flow cytometry to mark the differentiation trajectory similarly, we found that Progenitor (TCF1+) were dominantly associated with Q1 and Q2, putative ‘effectors’ (CD94+: CD94 or KLRD1 is the gene product of *Klrd1,* one of the markers of the T_EX_^KLR Eff^ subset) dominantly were Q2 and Q4 (with Q2>Q4) and ‘exhausted’ (PD1+CD38+) cells were enriched in Q4 and Q2 (Q4>Q2) (**Fig. 4d-f**), trends also recapitulated in spontaneous mammary PyMTChOVA tumors (**Fig. S7a-c**). The distribution of important transcription factor and other canonical marker expression also supported this broad relationship between T cell differentiation and CD69:TFP distributions (**Fig. 4g**). Transcripts for progenitor markers primarily concentrated in Q1, AP-1 factors in Q1 and Q2, while exhaustion-related markers largely occupied Q3 and Q4 states. Effector lineage and function-associated *Batf* and *Tbx21, Ifng* were highest in Q2 (Q2>Q4), along with signatures of non-canonical enhanced effectors derived from STAT5A overexpression(*37*) or MYB KO(*17*) (**Fig. 4g**). T_RM_ markers(*28*) *Prdm1,* and *Id2* tended to be higher in TFP^lo^ Q4 than TFP^hi^ Q2 in this context (**Fig. 4g**). A core tissue residency signature (*60*) also appeared split between Q2 and Q4 (Q2 > Q4) (**Fig. S6f**), while an orthogonal circulatory gene signature(*60*) was concentrated in Q1 and Q2 (**Fig. S6f**) – consistent with the early arrivals and those in the dLN being TFP^hi^. Identifying T_RM_-like cells by CD103 and CD69 expression by flow cytometry, we found their relative abundances in the Q2 and Q4 states to be similar (**Fig. S6g**), although the frequency of such cells among intratumoral OT-Is in this relatively short time of residence was only around 1% on average (**Fig. S6h**). A gene signature associated with transitory checkpoint blockade associated CD101-Tim3+ effectors were highest in Q2 and that of its CD101+Tim3+ terminally exhausted counterpart (*13*) was highest in Q4 followed by Q3 (**Fig. 4g**).

Within the entire population we found enhanced expression of intracellular cytokines TNF-α, IFN-γ and cytotoxicity-associated GzmB in intratumoral Q2 cells as compared to their Q4 counterparts, with restimulation post-sorting (**Fig. 4h-i, Fig. S7d**) as well as at baseline (**Fig. 4j, Fig. S7f, g**). Notably, Q1 cells were comparably cytokine and granzyme positive upon restimulation as Q2, while Q3 cells were similarly low response as Q4 (**Fig. 4i, Fig. S7d**). Without restimulation, Q2 cells were significantly more cytokine and granzyme producing that Q1 and the same was true for Q4 and Q3 (**Fig. S7e**), suggesting the conversion of re-activated cells from Q1 and Q3 to Q2 and Q4 respectively. The observation that effector-like and exhausted cells (delineated in single-cell transcriptomes and flow cytometry) both showed evidence of Q2 cells (**Fig. 4c, Fig. S6d**) implied the possibility that those demarcations may not fully capture the effector function potential of these cells. We thus interrogated Q2, as compared to Q4 cells from distinct differentiation states to determine whether there were variations in functional potency within each given classification. We gated further on subsets that were split between the Q2 and Q4 state – namely an early progenitor Ly108+CD69+(*11*) (56% Q2 and 44% Q4: **Fig. 4k**), an effector-like CD94+ (33% Q2 and 46% Q4: **Fig. 4l**) and terminal exhaustion-like PD1+CD38+ (23% Q2 and 53% Q4: **Fig. 4m**) subset, appearing at different junctures of the T_EX_ differentiation trajectory. Within Ly108+CD69+ (**Fig. 4k**) subset, cells in the Q2 state were higher than those in Q4 in TNF-α and IFN-γ (but not GzmB). Of note, a second progenitor population Ly108+CD69-(*11*), split between Q1 and Q3 showed lower cytokine and granzyme levels overall than Ly108+CD69+ and no significant difference between Q1 and Q3 (**Fig. S7h**). Further, within CD94+ (**Fig. 4l**) and PD1+CD38+ subsets as well (**Fig. 4m**), cells in the Q2 state consistently showed higher expression of these cytotoxic molecules in most cases than the Q4 counterparts (only TNF-α within CD94+ was not significantly different). The Ly108-CD69+ terminal subset aligned with the latter (PD1+CD38+) subset (∼83% of Ly108-CD69+ cells were PD1+CD38+: **Fig. S7i**), consistent with their association to terminal exhaustion, and therefore demonstrated similar Q2 vs. Q4 differences in cytokine and granzyme expression (**Fig. S7j**). Thus, this delineation of *Cd69* transcription (by TFP) among CD69+ cells provides an additional layer indicative of relative functionality of CD8 T cells over and above the established trajectory of an intratumoral T_EX_ differentiation landscape.

### Evidence of enhanced Q2 (TFP^hi^CD69^+^) CD8 T cell function during tumor regression

We next tested whether CD8 T cells in this activation state Q2 would be more prominent during a productive anti-tumor response such as T cell-mediated tumor control. To do this, we investigated subcutaneously injected MC38chOVA (MC38 colorectal cancer cell line modified with the same chOVA construct as PyMTchOVA (*54*) and B78chOVA) tumors that are actively controlled in response to the adoptive transfer of ovalbumin-specific OT-I T cells, whereas B78chOVA are not (**Fig. 5a**). We found that *Cd69*-TFP;OT-I T cells in regressing MC38chOVA tumors retain their predominantly Q2 (CD69^+^TFP^hi^) phenotype even at d12 post adoptive transfer, in contrast to those in the growing B78chOVA tumors (**Fig. 5b, Fig. S8a**). Conversely, the chronic activation-induced Q4 (CD69+TFP^lo^) subset was significantly lower in MC38chOVA compared to B78chOVA (**Fig. 5b**). However, the relatively high TFP^hi^ proportion (at least 60% at all time points) in the corresponding dLNs was similar in both (MC38chOVA :**Fig. S8b,** B78chOVA : **Fig. 2c, d**).

**Fig. 5:**
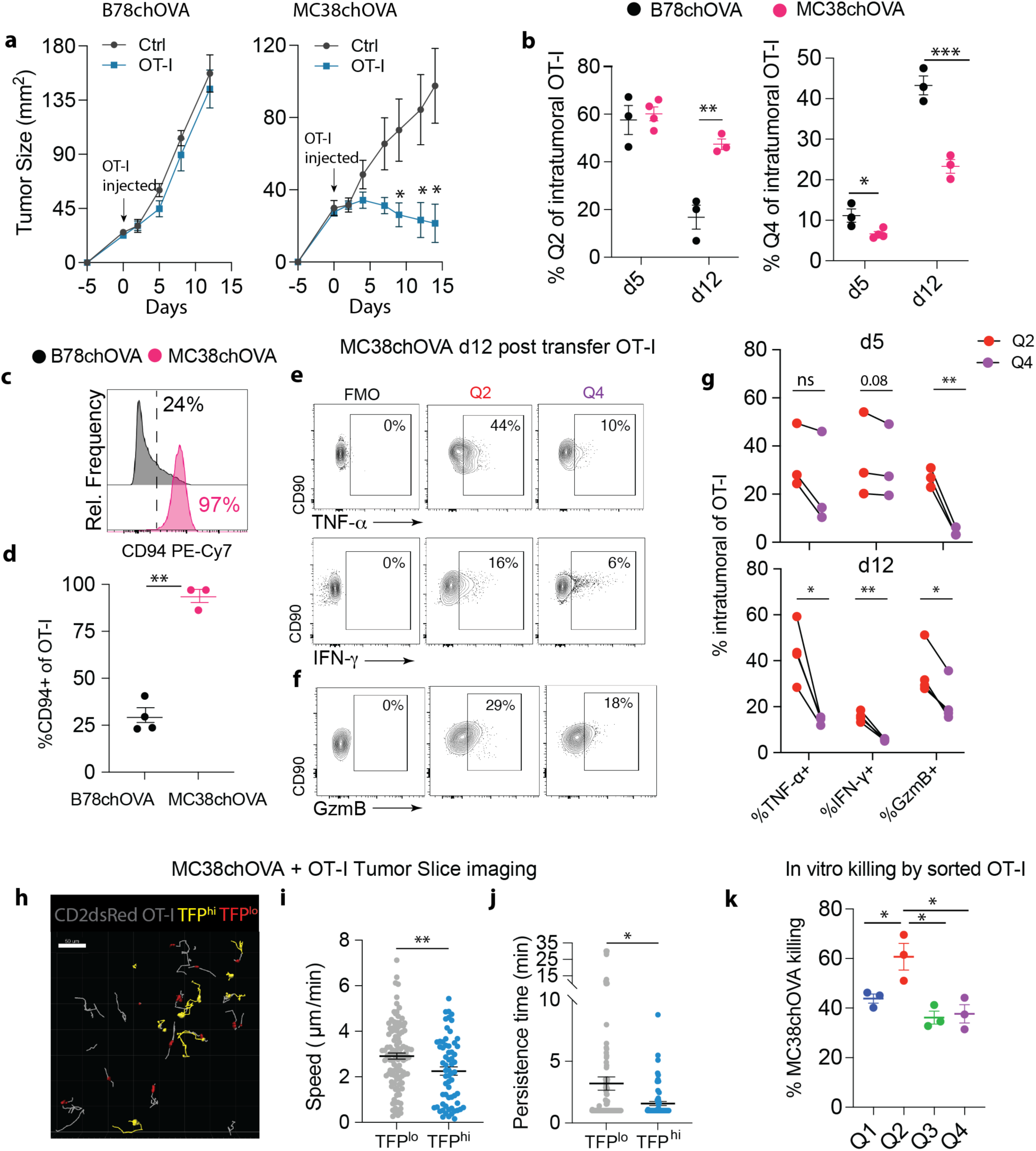
Evidence of TFP^hi^CD69+ (Q2) CD8 T cell function in a model of tumor regression: **(a)** Tumor growth curves of B78chOVA and MC38chOVA with and without OT-I transfer 5 days post tumor injection, as indicated by arrows (n=5-6/group); **(b)** quantification of the percentage of Q2 and Q4 cells at d5 and d12 post adoptive transfer in the two tumor models; **(c)** typical CD94 expression in Intratumoral OT-Is in B78chOVA and MC38chOVA tumors d12 post-adoptive transfer and **(d)** corresponding quantification; **(e-g)** Flow cytometry plots showing **(e)** intracellular cytokine and **(f)** GzmB expression in OT-I T cells in the Q2 vs. Q4 state d12 post adoptive transfer in MC38chOVA tumors and **(g)** quantification of the same at d5 and d12; **(h)** representative tumor slice image showing cell migration tracks associated with TFP^lo^ (red) and TFP^hi^ (yellow) CD2dsRed;OT-I T cells and corresponding quantification of **(i)** cell speed and **(j)** persistence of migration; **(k)** in vitro killing of MC38chOVA cells by OT-I T cells sorted by CD69:TFP quadrants from MC38chOVA tumors 8d after adoptive transfer; Bar plots show mean +/− SEM; null hypothesis testing by ratio paired t test (g), ANOVA with post-hoc Holm-Šídák test (b, k), unpaired t test (c) and Mann-Whitney test (i, j). *p<0.05, **p<0.01.

From d5 to d12, transferred OT-Is in MC38chOVA tumors showed a decline in Ly108 expression, and a concomitant increase in exhaustion-associated co-expression of PD1 and CD38, although the percentage of the latter was lower than that in B78chOVA (**Fig. S8c:** MC38hOVA**, Fig. S4b**: B78chOVA). Further investigation revealed the OT-I cells in the MC38chOVA tumors were ∼80% CD94 (KLRD1)+ at d5, which rose to be >90% at d12 (**Fig. S8c**). This was significantly higher compared to about 25% in the B78chOVA model (**Fig. 5c, d**), indicating a dominant KLR-effector phenotype in the MC38chOVA(*36*). Once again, the CD103+ subset of the transferred OT-Is was only around 1% (**Fig. S8c**), therefore ruling out T_RM_-like cells as the dominant potent effectors in this context. Among these largely KLRD1+ OT-I T cells in MC38chOVA, those in the Q2 state again tended to express markedly higher levels of TNF-α and IFN-γ and GzmB than Q4 counterparts, consistent with B78chOVA (**Fig. 5e-g**). These trends were already prevalent at d5 and showed a stronger divergence of phenotypes at d12 (**Fig. 5g**). Further evidence of in situ functional engagement of TFP^hi^ cells was obtained by two-photon microscopy where those cells could be identified by analysis of TFP levels over non-TFP controls (**Fig. S8d**). This analysis demonstrated enhanced cell arrest of the TFP^hi^ (majorly ∼75% Q2) cells within MC38chOVA tumor slices harboring adopted OT-Is, with lower speed (**Fig. 5h, i**), directional persistence (**Fig. 5j**), and overall motility (**Fig. S8e)**. In both mouse(*61*) and human(*57*) tumors, these traits are associated with lower levels of exhaustion. In addition, among the OT-I cells sorted from MC38chOVA tumors by CD69:TFP quadrants, those in the Q2 state also showed the highest killing capacity when exposed to MC38chOVA cells in vitro (**Fig. 5k**). These data substantiate the use of TFP and CD69 to mark a potent activation state of intratumoral CD8 T cells in mice.

### Demarcation of *Cd69* RNA levels enables functional delineation of CD69+ CD8 T cells across exhausted T cell subsets in human head and neck tumors

We next sought to independently identify similar CD8 T cell activation states in human tumors, now using multimodal CITE-Seq on Head and Neck Squamous Cell Carcinoma (HNSC) tumor biopsies (**Fig. 6a**). Within tumor-derived cell populations, we gated on CD8 T cells using antibody-tagged markers (**Fig. S9a**), and simultaneous readouts of Cd69 mRNA and surface CD69 protein expression allowed CD8 T cells to be gated into 4 quadrants (akin to Cd69-TFP reporter mice) (**Fig. S9b)**. Unbiased combined protein-RNA driven weighted nearest neighbor determination differentiated CD8 T cells into 7 major clusters and 2 minor, atypical clusters (MT+ and CXCL2+) (**Fig. 6b**). Based on the DEGs and overlaying exhaustion and naïve markers that were detected at the surface protein level by antibody-derived tags (**Fig. 6c**), we identified a naïve (Tn), a memory (Tem) and 5 exhaustion-associated clusters, including a proliferative and a CXCL13-expressing subset (**Fig. 6b, S9c, d**). In contrast to the Naïve and exhaustion scores, expression of progenitor and effector gene expression previously identified in chronic infection in mice(*12*) revealed diffuse expression in the major clusters (**Fig. S9e**). This may be expected since the trajectory of differentiation and typical subset identities in such human tumors are not identical to the T_EX_-differentiation trajectories in mice. Therefore, we used scSeq data corresponding to a recently published pan-cancer CD8 T cell atlas(*62*) and overlaid the DEGs for each major cluster in our dataset onto the established nomenclatures in the published atlas (**Fig. S9f, g**). This analysis supported the classification of the five exhausted (Tex) clusters as ‘early’, intermediate or ‘int’, ‘late’, cycling or ‘cyc’ and ‘CXCL13’, while Tn and Tem signatures corresponded directly to naïve and memory clusters in the atlas (**Fig. S9g**). Among the Tex clusters, the Tex^Early^ closely resembled TCF1+ Tex and OXPHOS-Tex from the atlas, the Tex^Int^ bore mild similarities to one NK-like Tex cluster, while Tex^Late^ and Tex^CXCL13^ were similar to Terminal Tex cluster in the atlas, which also expressed *CXCL13* (**Fig. S9c, g**). As in mice, Q2 (Cd69RNA^hi^CD69Protein^+^) and Q4 (Cd69RNA^lo^CD69Protein^+^) states in this human tumor sample also spanned several clusters (**Fig. 6d**). A majority of the exhausted CD8 T cells existed in the Q4 state, as indicated by the coincidence of the exhaustion score (based on protein expression) and the Q4 cells (**Fig. 6c, d**). Representation of the Q2 state was skewed towards the Tn, Tem and Tex^Early^ and that of Q4 towards the Tex^Late^, Tex^CXCL13^ subsets and the Tex^Int^ cluster (**Fig. 6e**).

**Fig. 6:**
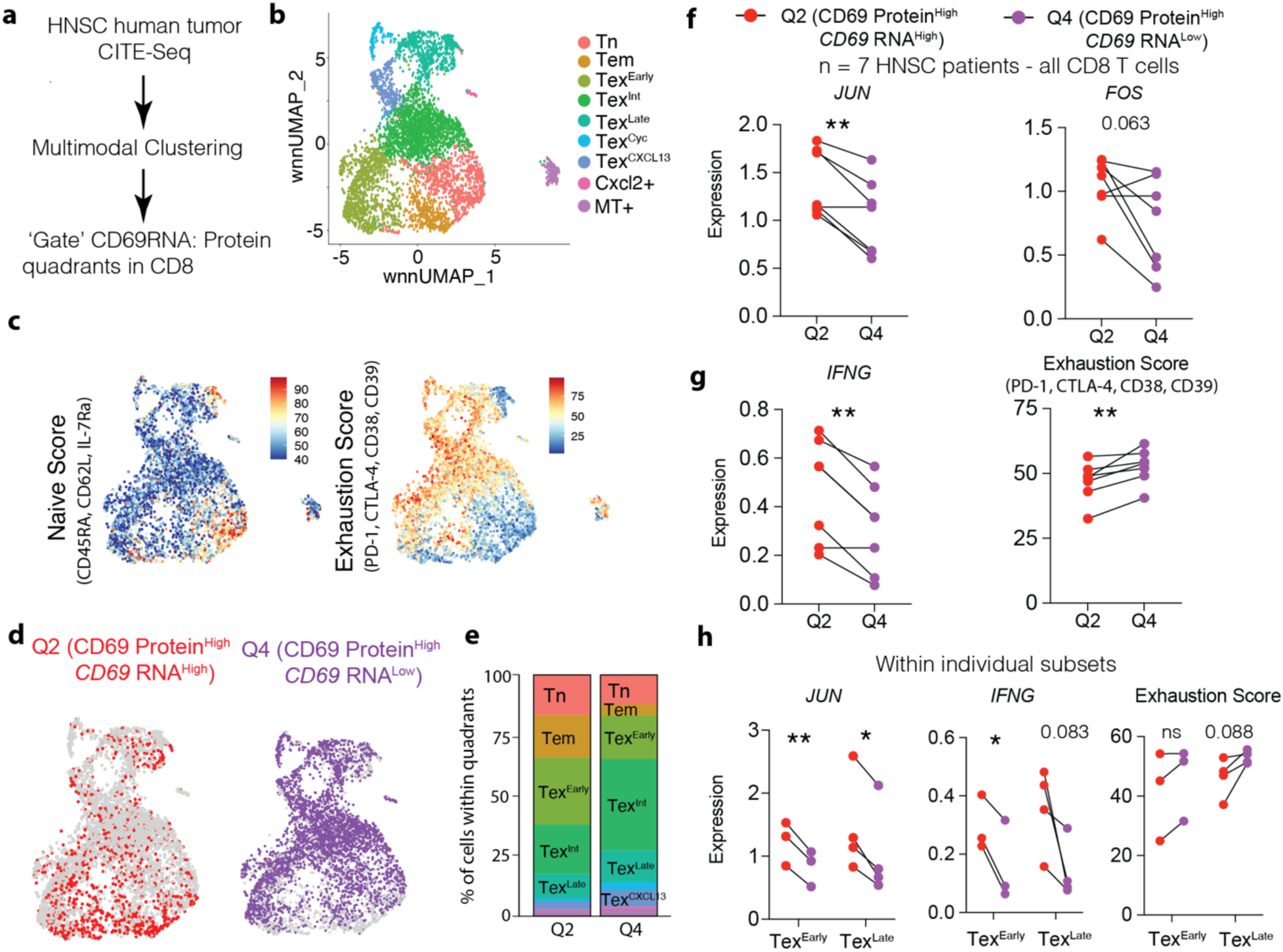
Simultaneous marking of Cd69 RNA and protein expression by CITE-Seq delineates intratumoral CD8 T cell activation states in head and neck squamous cell carcinoma: **(a)** Schematic description of human HNSC tumor CITE-Seq analysis; **(b)** UMAP showing weighted nearest neighbor (WNN) determined clusters by multimodal RNA and Protein analysis; **(c)** overlay of Naïve and Exhaustion score and **(d)** that of Q2 and Q4 cells (determined by gating on CD69 Protein and RNA) on the WNN-UMAP; **(e)** Stacked barplot showing relative contribution of the various subsets to the Q2 and Q4 states, as defined by Cd69 RNA and CD69 surface protein; Patient-by-patient expression of (f) *JUN* and *FOS,* (g) *IFNG* and Exhaustion markers in Q2 and Q4 CD8 T cells and (h) *JUN, IFNG,* Exhaustion markers between Q2 and Q4 within individual Tex^Early^ and Tex^Late^ subsets. *p<0.05, **p<0.01, ns = no significance by paired t tests (f-h).

Having classified both the Tex subsets and the Q1-Q4 quadrants within these intratumoral CD8 T cells, we then investigated the patterns of functional gene expression between the Q2 and Q4 states. Using the CITE-Seq data in a patient-by-patient analysis (**Fig. S9h**), we found that Q2 cells expressed significantly higher levels of AP-1 transcription factors *JUN* and *FOS* (STAT5A was poorly represented in this dataset) with 7 head and neck cancer patients as biological replicates (**Fig. 6f**). Enhanced expression of *IFNG* as well as a decreased Exhaustion score (calculated from antibody tags to exhaustion-related surface proteins) in Q2 as opposed to Q4 cells (**Fig. 6g**) indicated similar, functionally divergent activation states, as those elucidated in mice. Notably, as with the murine scSeq data, T_RM_-like cells delineated by CD103 protein expression were represented both in Q2 and Q4 states (**Fig. S9i**). We next interrogated whether the features of this delineation remained true within different subsets of cells, by delving into the individual Tex^Early^ and Tex^Late^ subsets, with sufficient representation (using a threshold of 50 cells in a given sub-group for obtaining reliable gene signature scores) in 3 and 4 patients respectively as biological replicates. Even within these distinct Tex subsets, the same trends of enhanced *JUN* and *IFNG* expression and decreased Exhaustion (only Exhaustion in Tex^Early^ remained indifferent) were observed (**Fig. 6g**), indicating a similar axis of effector function over and above the Tex differentiation trajectory.

### Delineation of potent effector states across human cancers

To the extent that transcriptomic signatures can capture the multimodal delineation offered by our reporter and then by CITE-Seq, these are useful in their imminent applicability to the already existing transcriptomic data on human tumors. With this objective, we obtained DEGs between Q2 and Q4 states from CITE-Seq. The Q2 signature included not only *CD69,* but also its upstream transcription factors *JUN, FOS* and *NR4A1*, while the Q4 signature largely comprised of actin-related genes and *CD74* (**Fig. 7a**). We then overlaid the expression of the gene signatures for Q2 and Q4 onto the pan-cancer CD8 T cell atlas described above(*62*) (**Fig. 7b**). We arranged the well-represented and characterized clusters (therefore excluding Tc17 (IL-17-related) and the NME-1+ clusters) in decreasing order of the naïve gene expression score (*SELL, IL7R, TCF7, LEF1, CCR7*) (**Fig. 7c**). As expected, the exhaustion gene expression score (gene list as previously described(*63*)) in the same clusters displayed approximately the opposite (increasing) order (**Fig. 7c**). Notably, while the Q4 score tended to coincide in pattern of expression with the exhaustion score, the Q2 score was distinct from both Naïve and Exhaustion score expressions (**Fig. 7c**). The Q2 score was prominent in the intermediate NK-like CD8 T cells expressing effector markers, while the Q4 signature associated highly with *GZMK* and *CXCL13* expressing T_EX_ subsets, which appear later in the predicted differentiation towards exhaustion (**Fig. 7c**).

**Fig. 7:**
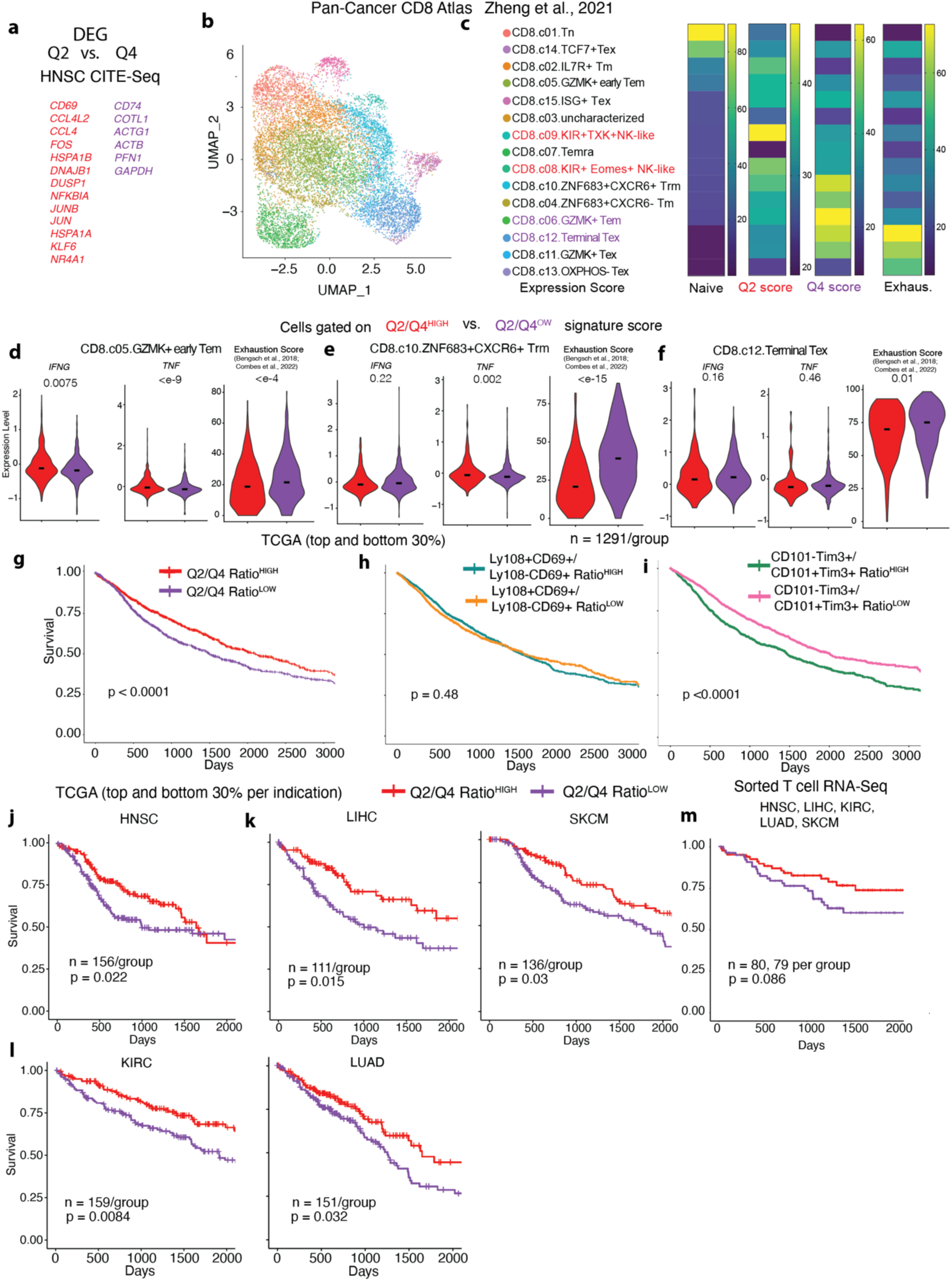
Transcriptomic signatures of Cd69RNA^hi^CD69Protein^hi^ (Q2) state associates with anti-tumor effector function in human cancers: **(a)** Differentially expressed genes of cells in Q2 vs. Q4 (|avg_logFC|>0.4, pval_adj <0.01) from the HNSC CITE-Seq data in Fig. 6 used to generate Q2 and Q4 signature scores; **(b)** UMAP representation of computationally-derived subsets among CD8 T cells in a pan-cancer T cell atlas(*62*), color-coded subset labels are as shown in (c); **(c)** Heatmap showing Naïve (*Sell, Il7r, Tcf7, Lef1, Ccr7*), Q2 (from a), Q4 (from a) and exhaustion(*63*) signature scores in the major CD8 T cell subsets from the same pan-cancer dataset, top two subsets in Q2 and Q4 score expression are labeled red and purple respectively, clusters are arranged in descending order of expression of the naïve score; expression of *IFNG*, *TNF* and Exhaustion signature score within the **(d)** Early Tem, **(e)** CXCR6+ Trm and **(f)** Terminal Tex clusters grouped by Q2/Q4^high^ and Q2/Q4^low^ (top and bottom 33% of the Q2 score/Q4 score ratio); **(g-i)** Kaplan-Meier survival curves from a quality controlled subset of TCGA data (*63*) corresponding to all indications stratified by the ratio of **(g)** Q2 and Q4 expression scores, **(h)** signatures corresponding to Ly108+CD69+ and Ly108-CD69+ subsets (Signature 1 and Signature 7 from (*11*)) and **(i)** CD101-Tim3+ effector and CD101+Tim3+ terminal subsets (*13*); Kaplan-Meier survival curves from a curated list of TCGA data(*63*) corresponding to **(j)** HNSC, **(k)** LIHC, SKM **(l)** KIRC and LUAD indications, patients stratified by the ratio of Q2 and Q4 expression scores; **(m)** Kaplan-Meier survival curves from a cohort of patients of combined HNSC, LIHC, SKM, KIRC and LUAD indications, with patients stratified by the ratio of Q2 and Q4 expression scores from bulk RNA-Seq of sorted T cells(*63*) number of patients per group and p-value for log-rank tests are noted in g-l and p-values for Wilcoxon test in d-f are noted. Black bars represent median in d-f.

As with the mouse scSeq, in this analysis of human tumor-infiltrated CD8 T cells, we further included previously identified exhausted T cell subsets in addition to our Cd69 RNA-based delineations. Specifically, transcriptomic signatures reported for phenotypically opposing Ly108+CD69+ vs. Ly108-CD69+(*11*), and CD101-Tim3+ vs. CD101+Tim3+ subsets(*13*) using human ortholog genes were examined. Cell-by-cell expression of Ly108-CD69+ and CD101+Tim3+ signatures (both of which are associated with terminal exhaustion) moderately correlated with each other (rho = 0.48) (**Fig. S10c**). The Q4 signature showed weaker (rho = 0.27, 0.25) correlation with both aforementioned signatures, while Q2 was largely uncorrelated with all other signatures (**Fig. S10c**). Notably these pairs of gene signatures (Q2 vs. Q4, Ly108+CD69+ vs. Ly108-CD69+ and CD101-Tim3+ vs. CD101+Tim3+ subsets) were not negatively correlated with their counterparts (**Fig. S10c**). Therefore, we gated on cells based on the ratio of Q2/Q4 signature scores within three distinct subsets - Gzmk+ early Tem, ZNF683+ T_RM_ and Terminal Tex - to test whether the Q2 and Q4 states represented divergent functional states within and across these subsets. Overall, the Q2/Q4 ratio^HIGH^ cells showed higher expression of effector cytokine transcripts *IFNG, TNF* and lower Exhaustion as compared to the Q2/Q4 ratio^LOW^ cells within these subsets, although the extent of these differences varied (**Fig. 7d-f**). In contrast, cells gated on the value of Ly108+CD69+/Ly108-CD69+ signature ratio were more often indifferent or skewed in the opposite direction with respect to the same metrics (**Fig. S10a**). The CD101-Tim3/CD101+Tim3+ ratio, while more variable showed some similarities with the trends in the Q2/Q4 delineation (**Fig. S10b**).

To interrogate how these Q2 and Q4 states associate with anti-tumor function in human tumors, we next used these gene signatures to query patient outcome data. We stratified HNSC patients from the cancer genome atlas (TCGA) using the same Q2 and Q4 signatures generated from HNSC CITE-Seq, into the top and bottom 30% based on the expression of the ratio of Q2/Q4 signature scores. HNSC patients with gene expression patterns skewed towards the Q2, as opposed to the Q4 state showed significantly better overall survival (**Fig. 7j**). In fact, patients scoring high on the Q2/Q4 ratio score showed significantly better overall survival in all indications combined (**Fig. 7g**). In contrast, the Ly108+CD69+/Ly108-CD69+ signature ratio was indifferent to (**Fig. 7h**) and the CD101-Tim3/CD101+Tim3+ ratio correlated negatively (**Fig. 7i**) with patient survival in the same TCGA cohort. While these results do not diminish the value of these previously described subsets for mapping T cell exhaustion, and indeed those signatures may be confounded by the presence of other cell types in their application to bulk TCGA RNAseq data, they do support the Cd69 RNA-based delineation as a distinct additional categorization of biological and clinical relevance.

By indication, in addition to HNSC, Q2/Q4 ratio was a significant survival prognostic in multiple indications including Hepatocellular (LIHC), Melanoma (SKCM), Kidney (KIRC) and Lung (LUAD) cancers (**Fig. 7k, l**), and in other indications (Bladder BLCA, Colorectal COAD, Glioblastoma GBM, Gynecological GYN, Pancreatic PAAD, and Sarcoma SRC) remained neutral to overall survival (**Fig. S10d, e**). To confirm that this association with survival was T cell-dependent, we next used a smaller bulk RNA-Seq dataset from sorted intratumoral conventional T cells that was part of a recent pan-cancer census(*63*). We applied the same strategy of stratifying patients (split at the median here to maximize number of samples) based on the Q2/Q4 ratio. Combining the above-mentioned indications (LIHC, SKCM, KIRC, LUAD, HNSC) where Q2/Q4 was associated with increased survival in TCGA, we found a clear trend towards a similar association even in this smaller dataset (∼80 patients/group) (**Fig. 7m**). Also consistent with the TCGA analysis, in this pan-cancer cohort, T cell specific Q2/Q4 ratio remained neutral to overall survival in other indications (BLCA, COAD, GBM, GYN, PAAD, SRC) combined (**Fig. S10f**). While these distinct outcome cohorts may be attributed to the heterogeneity across cancer types and potentially distinct underlying rate-determining steps for anti-tumor immunity, it is notable that in none of the indications this ratio emerges as a negative prognostic. Together, these data show that this delineation of CD69+ CD8 T cells by Cd69 RNA into potent and dysfunctional activation states is functionally relevant in many human cancers. Delineating this axis of potent effector states by tracking *Cd69* RNA provides an additional dimension to map the contours of the exhaustion-dominated T cell functional landscape in tumors.

## Discussion

In summary, we have defined a multimodal approach using transcript and protein levels of CD69 simultaneously to find potent effector CD8 T cell states over and above the exhausted T cell lineage in tumors. Our data suggests that such states may be naturally prominent in certain tumor microenvironments (MC38), and rare in others (B78/B16), with functional consequences. By determining functional states associated with patterns of *Cd69* RNA and protein expression, our data lend further relevance and context to the discordance between this transcript and its protein expression in T cells(*42, 43*). This agrees with the distinct kinetics of transcripts and proteins associated with many genes in the T and B cell repertoire (*44, 45*). Associated with tissue retention(*46, 56*) or stimulation in general, the relationship between CD69 protein expression and intratumoral T cell function is ambiguous. In contrast, we show that higher expression of *Cd69* RNA indicates a poised or potent activation state and therefore indicative of enhanced functional potential. The systematic use of *Cd69* transcription, along with its surface protein expression may be similarly applicable in other important contexts including vaccination, resident memory formation and autoimmunity, to directly identify and study potent activation states of lymphocytes in situ. While the fast maturation time of monomeric TFP allows concordant recording of new transcription, the longer half-life of the fluorescent protein makes it insensitive to rapid changes as a reporter. Rather, the decline in TFP expression should be thought of as a delayed reporting of decreasing transcription integrated over a period of days. Further, transcription and detectable transcript levels may also be different in some conditions where mRNA degradation(*43*) rather than translational repression dominates the post-transcriptional regulation. In this context, further studies using the inducible Cre in the reporter construct to lineage trace activated cell states marked by the reporter may be leveraged to further understand the nuance of this biology. Such timed Cre-mediated conversions may further allow marking of potent effector T cells for imaging as well as specific depletion in future studies.

Our results demonstrate that among CD69+ activated CD8 T cells, those with higher *Cd69* transcription, and associated higher levels of AP-1 transcription factor expression, are functionally distinct from those with lower levels of these transcripts. These data are consistent with separate previous reports showing the overexpression of AP-1 factors JUN, STAT5A and BATF in CD8 T cells leading to enhanced functional capacities(*37, 38*), (*39*). In the presence of AP-1 factors and NFAT, BATF can promote potent effector function and counteract exhaustion(*39*) – our data indicates that by the indirect recording of AP-1, NFAT activity by Cd69 transcription, this strategy illuminates those cells in which such functional signaling is active.

Accumulating evidence suggests that exhausted CD8 T cells in the TME continue to receive sub-optimal TCR stimulation, keeping them locked in a dysfunctional activation state(*21, 64*). Within and across broadly defined subsets, the delineation of activation states here serves to further refine our understanding of the anti-tumor potential of intratumoral CD8 T cells. Our data supports the delineation of Q2 (RNA^hi^Protein^+^) and Q4 (RNA^lo^Protein^+^) as ‘potent’ and ‘chronic’ activation states respectively. Q1 (RNA^hi^Protein^-^) on the other hand represents a ‘poised’ state, while Q3 (RNA^lo^Protein^-^) is a quiescent state encompassing both certain naïve and bystander cells, as well as inactive exhausted cells. In this way, these results should be understood to not be ‘in place of’ but ‘in addition to’ the already existing mapping of exhausted T cell states which we have extensively referenced and drawn from in this work. The application of this methodology in the CITE-Seq context demonstrates the potential to find functionally relevant activation states in human cancers. As such, the gene signatures for the divergent activation states derived from head and neck tumors may be further refined to be robust across species and indications – perhaps with the aid of bulk RNA-Seq or ATAC-Seq modalities, which may offer a deeper look into such states.

The use of CD69 protein as a primary marker of specific T cell states is a point of consideration for future studies. Our data show that repetitive TCR stimulation and exposure to antigen-bearing TME are sufficient for the drop in relative transcription levels and TFP and the resultant functional differences between TFP^hi^ and TFP^lo^ subsets. However, given other stimuli that can induce CD69 expression, it is not necessary that Q2 and Q4 cells be produced by TCR-dependent mechanisms. It remains to be explored whether other forms of repetitive stimulation, or successive recall (in the case of memory cells) can drive such a feedback response associated with decreasing function. Overall, the interpretation of CD69 protein expression likely depends on these additional measures of feedback, such as its transcript levels which are shown here to be related to history and effector function.

Successful boosting of anti-tumor immunity in rare pockets of reactive immunity(*1, 65, 66*) may lead to the generation of stronger effector CD8 T cells, inducing functional features within subsets of exhausted T cells, perhaps without entirely reversing the differentiation favoring exhaustion. Indeed, atypical effector states have been amplified by the specific deletion or overexpression of exhaustion lineage-defining transcription factors(*17, 19*). These and other data(*36*) suggest that the T_EX_ subsets are not entirely devoid of plasticity. It is likely that T cells in various stages of T_EX_ differentiation within tumors may encounter distinct microenvironmental signals that push them towards an enhanced or diminished functional state. While the overarching trajectory of CD8 T cell differentiation in tumors may be set by exhaustion-lineage defining factors, there appear to be local and indeed global peaks and valleys of function that may be determined by additional axes of potency.

As exploration of spatial niches of functional immunity continue to drive the field, we posit that this approach to directly mark the potent effectors in the TME may be an important anchoring tool. Detecting, studying and ultimately enhancing functional anti-tumor CD8 T cells will lead to the discovery of novel strategies to drive better patient outcomes.

## Funding

National Institutes of Health Grants: NIH R01CA197363 and NIH R37AI052116

AR was supported by a Cancer Research Institute Postdoctoral Fellowship (CRI2940)

KHH was supported by an American Cancer Society and Jean Perkins Foundation Postdoctoral Fellowship

GH was supported by the National Science Foundation Graduate Research Fellowship Program (NSF2034836)

IZL was supported by Emerson Collective Health Research Scholars Program

We thank Dr. Emily Flynn, UCSF for guidance with the CITE-Seq analysis. We thank Dr. Jeroen Roose, UCSF, Dr. Julie Zikherman, UCSF and Dr. Susan Schwab, NYU for discussions and guidance. We thank Dr. Kelly Kersten, Sanford-Burnham Prebys, Dr. Mike Kuhns, University of Arizona and Dr. Miguel Reina-Campos, University of California San Diego, for critical reading of the manuscript. We also thank Dr. Reina-Campos for sharing data from Zheng et.al in an analysis-ready format. We thank members of the Krummel lab for their inputs to the manuscript.

## Author Contributions

Conceptualization: AR, MFK

Experimentation: AR, MB, GH, LFP, IZL

Mouse scSeq: KHH, AR

Human tumor data analysis: AR, AJC, BS

Human tumor data collection: VD, BD, AJC

Funding acquisition: MFK, AR, KHH

Writing: AR, MFK

Supervision: MFK

## Competing Interests

The authors declare no competing interests

## Data materials availability

Relevant data will be made publicly available before publication in its final form. Meanwhile, data will be available upon reasonable request, please contact the authors directly.

## Supplementary Materials

### Materials and Methods

#### Mice

All mice were treated in accordance with the regulatory standards of the National Institutes of Health and American Association of Laboratory Animal Care and were approved by the UCSF Institution of Animal Care and Use Committee. Cd69-TFP-CreER^T2^ (denoted as Cd69-TFP) mice in the C57BL6/J background were custom-generated from Biocytogen Inc. and then maintained heterozygous (bred to C57BL6/J wild type mice) at the UCSF Animal Barrier facility under specific pathogen-free conditions. C57BL6/J (wild type; WT), C57BL6/J CD45.1 (B6.SJL-Ptprc^a^ Pepc^b^/BoyJ), OT-I (C57BL/6-Tg(TcraTcrb)1100Mjb/J) mice were purchased for use from Jackson Laboratories and maintained in the same facility in the C57BL6/J background. For adoptive transfer experiments, CD45.1^het^; OT-I^het^; Cd69-TFP^het^ (denoted simply as CD45.1; OT1; Cd69-TFP) mice were used. Mice of either sex ranging in age from 6 to 14 weeks were used for experimentation. For experiments using the transgenic PyMTchOVA strain(*54*), mammary tumor-bearing females in the age range of 15 to 24 weeks were used. Adoptive transfer of T cells in these mice were done when mice developed at least 2 palpable tumors (> 25-30mm^2^).

#### Mouse tumor digestion and flow cytometry

Tumors from mice were processed to generate single cell suspensions as described previously(*67*). Briefly, tumors were isolated and mechanically minced on ice using razor blades, followed by enzymatic digestion with 200 μg/mL DNAse (Sigma-Aldrich), 100U/mL Collagenase I (Worthington Biochemical) and 500U/mL Collagenase IV (Worthington Biochemical) for 30 min at 37°C while shaking. Digestion was quenched by adding excess 1X PBS, filtered through a 100μm mesh, spun down and red blood cells were removed by incubating with RBC lysis buffer (155 mM NH_4_Cl, 12 mM NaHCO_3_, 0.1 mM EDTA) at room temperature for 10 mins. The lysis was quenched with excess 1X PBS, spun down and resuspended in FACS buffer (2mM EDTA + 1% FCS in 1X PBS) to obtain single cell suspensions. Similarly, tumor draining lymph nodes (dLN) were isolated and mashed over 100μm filters in PBS to generate single cell suspensions.

For each sample, 2.5-3 million cells/sample were stained in a total of 50µL of antibody mixture for flow cytometry. Cells were washed with PBS prior to staining with Zombie NIR Fixable live/dead dye (1:500) (Biolegend) for 20 min at 4°C. Cells were washed in FACS buffer followed by surface staining for 30 min at 4°C with directly conjugated antibodies diluted in FACS buffer containing 1:100 anti-CD16/32 (Fc block; BioXCell) to block non-specific binding. Antibody dilutions ranged from 1:100-1:400, optimized separately. After surface staining, cells were washed again with FACS buffer. For intracellular staining, cells were fixed for 20 min at 4°C using the IC Fixation Buffer (BD Biosciences) and washed in permeabilization buffer from the FoxP3 Fix/Perm Kit (BD Biosciences). Antibodies against intracellular targets were diluted in permeabilization buffer containing 1:100 Fc Block and cells were incubated for 30 min at 4°C followed by another wash prior to readout on a BD LSRII or Fortessa Cytometer.

#### Processing and flow cytometry analysis of other mouse organs

To phenotype T cells under from lymphoid organs homeostasis, spleen and inguinal, mesenteric and brachial lymph nodes were isolated and mashed over 100µm filters washed with 1X PBS to generate single cell suspension of lymphocytes. For splenic suspensions, RBC lysis was performed as described above before staining for flow cytometry.

To profile thymocytes, thymus was isolated, cut into small pieces with a razor blade and minced by using gentleMACS dissociator (Miltenyi Biotec) in RPMI. Next, the mixture was spun down and resuspended in the digestion mixture described above and allowed to digest with shaking at 37°C for 20 mins, following which, the remaining tissue was either minced again using the gentleMACS dissociator and/or directly mashed over a 100µm filter in FACS buffer to generate a single cell suspension, ready to be processed for staining and flow cytometry.

Skin digestion was done as previously described(*68*). Briefly, mice are shaved and depilated prior to removal of dorsal skin. The skin was then rid of fat, minced with scissors and razor blade in the presence of 1 ml of digest media (2 mg/ml collagenase IV (Roche), 1 mg/ml hyaluronidase (Worthington), 0.1 mg/ml DNase I (Roche) in RPMI-1640 (GIBCO). The minced skin was then moved to a 50 ml conical with 5 ml additional digest solution and incubated at 37°C for 45 min with shaking and intermittent vortexing before being washed and passed through a 70μm strainer prior to staining. TFP high vs. low gates were drawn by using a side-by-side WT control or using endogenous CD8 T cells in the context of adoptive transfer into a tumor-bearing mouse.

#### Tumor injections and adoptive transfer of CD8 T cells into tumors

The B78chOVA and MC38chOVA cancer cell lines, as previously described(*23, 67*), were generated by incorporating the same mcherry-OVA construct used to establish the PyMTchOVA spontaneous mouse line(*54*). For tumor injections, the corresponding cells were grown to near confluency (cultured in DMEM with 10% FCS (Benchmark) and 1% PSG (Gibco)) and harvested using 0.05% Trypsin-EDTA (Gibco) and washed 3x with PBS (Gibco). The number of cells to be injected per mouse was resuspended in PBS and mixed in a 1:1 ratio with Growth Factor Reduced Matrigel (Corning) to a final volume of 50μL per injection. The mixture was injected subcutaneously into the flanks of anesthetized and shaved mice. Tumors were allowed to grow for 14–21 days unless otherwise noted, before tumors and tumor-draining lymph nodes were harvested for analysis. CD8 T cells were isolated from CD45.1;OT-1;Cd69-TFP mice using the EasySep Negative Selection Kit (Stem Cell Bio), resuspended in 1X PBS at 10X concentration 100µL was injected into each tumor-bearing mice. For B78chOVA and PyMTchOVA tumors, 1 million and for MC38chOVA tumors, 200,000 CD8 T cells were injected retro-orbitally into each mouse either 5d (B78chOVA and MC38chOVA) post tumor injection or when mice had at least 2 palpable tumors (PyMTchOVA). Tumor measurements were done by measuring the longest dimension (length) and approximately perpendicular dimension (width) using digital calipers, rounded to one decimal place each.

#### Contralateral tumor injection and vaccination

5 days post B78chOVA tumor injection, equal numbers (1 million) CD8 T cells from a CD45.1;OT-I;Cd69TFP and P14;Cd69TFP mice were injected retroorbitally into each mice. Next day, gp33-41 subcutaneous peptide (Anaspec) vaccination was injected contralaterally to the tumor, with 50µg peptide + 50µL Common Freund’s Adjuvant (CFA, Sigma) along with 50µL PBS for a total volume of 100µL. The vaccination site was identified by a white, hardened subcutaneous mass and isolated and processed similarly to the tumor for flow cytometry.

#### In vitro stimulation of naïve CD8 T cells

CD8 T cells were isolated from Cd69-TFP or WT mice as described above and plated in a 96 well round bottom plate (Corning) at 80,000 cells/well in T cell media-RPMI (Gibco) + 10% FCS (Benchmark) + Penicillin/Streptomycin + Glutamine (Gibco). TCR stimulation was induced by adding anti-CD3/CD28 Dynabeads (Applied Biosystems) at the concentration of 2µL per 80,000 cells (1:1 ratio of cells:beads), the plate was briefly spun down to bring cells and beads together before incubation at 37°C for varying lengths of time. 55µM β-mercaptoethanol (BME; Gibco) was added to the T cell media during stimulation. For repeated stimulation assays, 2 wells of each sample at every time point were pooled for mRNA isolation and qRT-PCR, while 2 other wells were used as duplicates for flow cytometry. After each cycle, beads taken off each well and replated for resting in T cell media containing 10 U/mL of Interleukin-2 (IL-2; Peprotech). To restart each stimulation cycle, cells from each biological replicate were pooled, counted and Dynabeads were added at the appropriate concentration for a 1:1 ratio and redistributed into wells for incubation. To assay shorter stimulation times (3, 6, 18h), the same repeated stimulation and rest experiment was carried out but analyzing cells by flow cytometry at these early stimulation time points at each cycle.

For cytokine production assays, cells at the beginning of Cycle 3 were sorted by TFPhi vs. lo levels, with the non-reporter expressing control cells used to set the gate. Sorted cells were plated in a 96-well V-bottom plate either in T cell media or T cell media containing PMA (50 ng/mL; Sigma-Aldrich), Ionomycin (500ng/mL; Invitrogen) + Brefeldin A (3µg/mL; Sigma-Aldrich) and BME (Gibco) for 3h, before cells were collected for surface and intracellular staining for cytokines and granzyme B.

#### Sorting and qPCR, resting or restimulation of homeostatic CD8 T cells

To sort sufficient CD8 T cells from homeostatic lymphoid organs, CD8 T cells were first isolated from spleens and inguinal, brachial, mesenteric lymph nodes Cd69-TFP or WT mice using the EasySep Negative selection kit. These cells were then sorted on TFP^hi^ (top 15%), TFP^mid^ (middle 30%) and TFP^lo^ (bottom 15%) from each mouse separately and rested in T cell media containing 10 U/mL Interleukin-7 in a 96 well round bottom plate and assayed at 0, 24 and 48h. Likewise, for qPCR analysis of populations high, mid and low for TFP, these populations were sorted into cold T cell media, pelleted and subjected to RNA extraction and qPCR with primers for Cd69 and 18s rRNA as the reference gene. For the sort and restimulation experiment, Memory (CD44+CD62L+) TFP^hi^ cells and Effector (CD44+CD62L-) TFP^lo^ cells were sorted and incubated in T cell media + 55µM BME containing 1:1 anti CD3/CD28 Dynabeads in a 96 well round bottom plate with either 5µg/mL Actinomycin D (Sigma) in DMSO or DMSO alone (vehicle) for 3h, before profiling by flow cytometry. De novo CD69 surface expression was measured by the difference of CD69 MFI between the vehicle and Actinomycin D treated groups.

#### Restimulation and cytokine production of intratumoral CD8 T cells

OT-I T cells from B78chOVA tumors were sorted on a BD FACSAria Fusion or BD FACSAria2 (BD Biosciences) at d11-d13 post adoptive transfer of CD8 T cells from CD45.1; OT-I; Cd69-TFP mice, as described above. To prepare CD45-enriched fractions(*69*), tumors were digested as described above into single cell suspensions, centrifuged and resuspended in 30mL room temperature (RT) RPMI 1640. Then, 10mL Ficoll-Premium 1.084 (Cytiva) was carefully underlaid and the tubes centrifuged at 1025g for 20 mins at RT without braking. The resulting interface-localized cells were pipetted out, diluted in equal volume RPMI and centrifuged at 650g for 5 mins to collect the cells. This constituted a CD45-enriched fraction which was then processed for staining and FACS. The four CD69:TFP quadrants were sorted from each tumor sample (cells from 2-3 tumor samples were pooled for a single biological replicate) into serum-coated microcentrifuge tubes containing cold T cell media. These were subsequently plated in a 96-well V-bottom plate either in T cell media or T cell media containing PMA (50 ng/mL; Sigma-Aldrich), Ionomycin (500ng/mL; Invitrogen) + Brefeldin A (3µg/mL; Sigma-Aldrich) and BME (Gibco) for 3h, before cells were collected for surface and intracellular staining for cytokines and granzyme B.

To determine the conversion trajectory of Cd69:TFP quadrants, intratumoral OT-I T cells were similarly sorted by Cd69:TFP gating, and restimulated with excess anti CD3/CD28 Dynabeads in a 96 well V-bottom plate for 3d. Due to dearth of Q1 cells in the tumors at d12, we sorted those cells from the dLN instead. The resulting re-stimulated cells were assayed by flow cytometry to determine converted Cd69:TFP quadrant distributions from each group.

#### Long-term ex vivo tumor slice overlay

Tumor slice overlay cultures were adapted, modified and extended from previous work(*57*). For tumor slice overlay cultures, B78chOVA tumors were injected bilateral subcutaneously into the flank of anesthetized and shaved mice. Tumors were allowed to grow for 11 – 13 days. 96 hours prior to tumor harvest and slicing, CD8 T cells were isolated from CD45.1;OT-1;Cd69-TFP mice, as described above. Isolated CD8 T cells were activated via 1:1 culture with Dynabeads Mouse T-Activator CD3/CD28 (Invitrogen) in T cell media + BME in 96 well U-bottom plates for 48 hours. After activation, T cells were removed from Dynabeads rested in T cell media with supplemented 10 U/mL IL-2 (PeproTech) for 48 hours before use. For gating TFP high vs. low cells, CD8 T cells from CD45.1; OT-I mice were subjected to similar pre-treatment and profiled by flow cytometry side-by-side along with the CD45.1;OT-1;Cd69-TFP CD8 T cells at d0.

For slicing, tumors were harvested and stored in cold RPMI until use. Each well of a 24 well plate was pre-filled with cold RPMI and stored on ice. Tumors were embedded in 1.5-2% agarose gel, allowed to solidify, and sliced at a thickness of 350 – 400µm using a Compresstome VF310-0Z Vibrating Microtome (Precisionary). Slices were immediately stored in pre-filled 24 well plate on ice until use.

For the slice overlay, each well of a 24 well plate was pre-coated with 30µL of 1 part culture medium:4 parts Matrigel and allowed to solidify at 37°C. Tumor slices were removed from RPMI and excess agarose was trimmed from slice edges (leaving a thin halo of agarose around slices to use for handling). Slices were spread across solidified Matrigel bed in 24 well plates. Rested T cells were stained with Violet Proliferation Dye 450 (BD Biosciences) diluted 1:1000 in PBS at 10 × 10^6^ cells/mL for 15 minutes at 37°C. Cells were washed 2x with PBS and resuspended in T cell culture medium at 150 – 200 × 10^6^ cells/mL. 5µL cell suspension (0.5 – 1 × 10^6^ cells) was added directly on top of each slice and incubated at 37°C for 3 hours, with 5µL fresh media added to each slice every 30 minutes to prevent slices from drying out. After incubation, 30µL non-diluted Matrigel was added directly atop each slice and allowed to solidify at 37°C. 2mL T cell culture medium containing BME was added to each well. 1mL culture medium was removed and replaced with fresh medium every 24 hours throughout the experiment.

#### Single cell RNA Sequencing and Analysis

Adoptively transferred CD45.1; OT-I; Cd69-TFP CD8 T cells were sorted from B78chOVA tumors d12 post transfer into four populations based on the CD69:TFP quadrants (Q1: TFP+/CD69-, Q2: TFP+/CD69+, Q3: TFP-/CD69-, and Q4: TFP-/CD69+). Cells were sorted from 8 tumors grown in separate mice and pooled by quadrants for barcoding and analysis. Sorted cells were separately labeled with lipid and cholesterol-modified oligonucleotides (LMO’s) according to McGinnis et. al(*70*). Following 2 washes with PBS + 0.1% BSA, cells were pooled for encapsulation in one lane of a 10X 3’ v3 kit with a target cell number of 18,000.

Following construction of the GEX library (according to manufacturer’s instructions) and the LMO library(*70*), libraries were pooled at a 10:1 molar ratio for sequencing on the NovaSeq 6000. This resulted in 807M cDNA reads and 163M LMO reads. Transcript and LMO reads were counted using the CellRanger count function against the GRCm38 reference genome to generate feature barcode matrices. These matrices were loaded into Seurat and filtered to remove high mitochondrial % cells (> 15%) and cells with low nGene (< 200 genes). Cells were then demultiplexed using their LMO counts with cells having too few LMO nUMI or ambiguous identity (possible multiplets) filtered out using the demultiplex package(*70*). The resulting object had an average cDNA nUMI per cell of 7662 reads and average nGene per cell of 2115 genes and average LMO nUMI per cell of 1080 reads. The final object underwent scaling and then scoring for cell cycle signatures (S and G2M scores as computed using Seurat’s built-in CellCycleScoring function. The object then underwent regression for cell cycle effects (S and G2M score as described in the Seurat vignette) and percent mitochondrial reads before PCA. K-Means clustering and UMAP dimensional reduction was then performed on the first 16 PC’s.

To compare gene expression signatures of previously established subsets to the clusters in our dataset, we obtained the top 10 differentially expressed genes (sorted by log fold change and having an adjusted p-value <0.01) of T_EX_ clusters described in Daniel et al(*12*). Then we calculated gene expression scores corresponding to those subsets and mapped them onto the clusters in our data.

#### qRT-PCR

At designated time points, CD8 T cells were isolated from the 96 well culture plates, or CD8 T cells were sorted into T cell media and centrifuged. The supernatant was aspirated out and the pellets stored at −80°C until mRNA extraction using the RNEasy Micro Kit (Qiagen). Corresponding cDNA was synthesized from the mRNA samples using the cDNA amplification kit (Applied Biosystems). qPCR using pre-designed *Cd69* and *18s* probes (Invitrogen) with a TaqMan-based assay system (BioRad) or custom-made primers (iDT Technologies) for *Jun* (Fwd: 5′ ACGACCTTCTACGACGATGC 3′, Rev: 5′ CCAGGTTCAAGGTCATGCTC 3′)(*71*) and *18s* (Fwd: 5’ CTTAGAGGGACAAGTGGCG 3’, Rev: 5’ ACGCTGAGCCAGTCAGTGTA 3’)(*72*) using the SsoFast assay system (BioRad) was used to quantify transcripts in a BioRad CFX94 machine.

#### Human tumor samples

All tumor samples were collected with patient consent after surgical resection under a UCSF IRB approved protocol (UCSF IRB# 20-31740), as described previously(*63*). In brief, freshly resected samples transported in ice-cold DPBS or Leibovitz’s L-15 medium before digestion and processing to generate a single-cell suspension. The following cancer indications were included in the cohort: Bladder cancer (BLAD), colorectal cancer (CRC), glioblastoma multiforme (GBM), gynecological cancers (GYN), hepatocellular cancers (HEP), head and neck cancer (HNSC), kidney cancer (KID), lung cancer (LUNG), melanoma (MEL), pancreatic ductal adenocarcinoma (PAAD), sarcoma (SRC).

#### Transcriptomic analysis of human tumors

All tumor samples were collected under the UCSF Immunoprofiler project as described(*63*). Briefly, tumor samples were thoroughly minced with surgical scissors and transferred to GentleMACs C Tubes containing 800 U/ml Collagenase IV and 0.1 mg/ml DNase I in L-15/2% FCS per 0.3 g tissue. GentleMACs C Tubes were then installed onto the GentleMACs Octo Dissociator (Miltenyi Biotec) and incubated for 20 min (lymph node) or 35 min (tumor) according to the manufacturer’s instructions. Samples were then quenched with 15 mL of sort buffer (PBS/2% FCS/2mM EDTA), filtered through 100μm filters and spun down. Red blood cell lysis was performed with 175 mM ammonium chloride, if needed. Freshly digested tumor samples were sorted by FACS into conventional T cell, Treg, Myeloid, tumor and in some cases, stromal compartments and bulk RNA-seq was performed on sorted cell fractions. mRNA was isolated from sorted fractions and libraries were prepared using Illumina Nextera XT DNA Library Prep kit. The libraries were sequenced using 100bp paired end sequencing on HiSeq4000. The sequencing reads we aligned to the Ensembl GRCh38.85 transcriptome build using STAR(*73*) and gene expression was computed using RSEM(*74*). Sequencing quality was evaluated by in-house the EHK score, where each sample was assigned a score of 0 through 10 based on the number of EHK genes that were expressed above a precalculated minimum threshold. The threshold was learned from our data by examining the expression distributions of EHK genes and validated using the corresponding distributions in TCGA. A score of 10 represented the highest quality data where 10 out of 10 EHK genes are expressed above the minimum threshold. The samples used for survival analysis and other gene expression analyses had an EHK score of greater than 7 to ensure data quality. Ensemble gene signatures scores were calculated by converting the expression of each gene in the signature to a percentile rank among all genes and then determining the mean rank of all the genes in the signature.

#### Survival analysis

Survival analyses using the TCGA dataset was performed using the TCGA sub-cohort described in (*63*). Briefly, tumor RNAseq counts and TPM along with curated clinical data for 13 cancer types (BLCA, COAD, GBM, GYN (grouping OV, UCEC and UCS), HNSC, KIRC, LIHC, LUAD, PAAD, SARC and SKCM) was filtered down to include primary solid tumors and metastatic samples only. This reduced the TCGA sample set to 4341 tumor samples. Q2 and Q4 gene scores (refer to the CITE-Seq section for details on the generation of the corresponding gene lists) were generated by first normalizing (using percentiles) the expression values of each gene composing the signature across all patients, followed by averaging these normalized values for each patient. We then calculated the ratio of Q2/Q4 gene scores for each patient. For survival analysis, patients were split into either or (Q2:Q4 gene signature ratio)^HIGH^ vs (Q2:Q4 gene signature ratio)^LOW^ (top/bottom 30% respectively, n=1291) and analyzed using a log-rank test. Similarly, the same Q2:Q4 ratio was used to stratify patients from the recent pan-cancer cohort with sorted T cell bulk RNA-Seq (*63*). Here, patients from the indications where Q2/Q4 ratio showed a prognostic benefit in the TCGA were combined together and those that did not were combined separately. Within each of these groups, T cell specific Q2/Q4 signature score ratio high and low were defined simply by the median of this ratio, and the survival of patient groups stratified in this way were tested by the log-rank test.

#### Analysis from published datasets

Available, curated RNA-Seq data (*8, 11, 23*) on *Cd69* and upstream transcription factor expression was plotted directly without modification.

A curated R object derived from Zheng et al.(*62*) generously shared by Dr. Miguel Reina-Campos, UCSD, was used for analysis. Ensemble gene signatures were scored as mentioned above and plotted onto pre-existing UMAP dimensional reduction and already annotated cell clusters. While exhaustion genes were obtained from previously published work(*63*), *TCF7, SELL, LEF1, CCR7, IL7R* genes were used for the Naïve score. Q2 and Q4 expression scores (refer to the CITE-Seq section for details on the generation of the corresponding gene lists) were generated by the percentile scoring method as described above. Likewise, Ly108+CD69+ and Ly108+CD69-gene signatures were generated from gene lists corresponding to Signature 1 and Signature 7 from Beltra et al. 2020 (*11*) and the CD101-Tim3+ and CD101+Tim3+ gene lists were obtained from Hudson et al. (*13*) by filtering the top 25 by log fold change with p values <0.01. From these murine gene lists, some genes that did not have clear human orthologs or showed no expression in the Pan-cancer atlas dataset in question were excluded. Cells in the pan-cancer CD8 atlas were then gated by the ratio of Q2/Q4, Ly108+CD69+/Ly108-CD69+ and CD101-Tim3+/CD101+Tim3+ expression ratios (top and bottom 33% of the ratios) and queried for the expression of cytokine and exhaustion-related markers.

The signatures of Myb cKO T_EX_ cells(*17*) and STAT5A over-expressed CD8 T cells(*37*) were derived by the same method using the top10 differentially expressed genes (by log fold change, with an adjusted p-value threshold <0.01). CD101-Tim3+, CD101+Tim3+ signatures were generated from published gene lists (*13*) as described above and the same procedure was followed for the circulating vs. resident signatures (*60*), in the latter taking the top 20 genes by log fold change and filtered by p-value <0.01. These were applied directly to the murine scSeq data sorted by Cd69:TFP quadrants.

#### In vitro Killing Assay

MC38chOVA tumors with adoptively transferred Cd69-TFP-OT-I CD8 T cells in WT B6 mice were harvested at d8 post T cell transfer, digested as mentioned above, and sorted by CD69: TFP quadrants into cold T cell media. Sorted cells were centrifuged, resuspended in fresh, warm T cell media with BME and added onto MC38chOVA cells plated ∼24h prior in flat-bottom 96 well plates. To each well containing 5000 MC38chOVA plated 24h prior, 5000 sorted T cells were added. As with the sort and restimulation experiments, each such collection of 5000 cells from a particular quadrant was pooled from 2-3 tumors and treated as a single biological replicate. Each experiment involved 7-8 tumors to obtain at least 3 biological replicates. Technical replicates were included and averaged wherever possible, i.e., at least 10,000 cells were sorted from a given quadrant and biological replicate. Percentage killing was obtained by measuring the fractional loss of live cells at 36h in no T cell vs. T cell added conditions relative to 0h. Live cell numbers from each condition was accurately measured by lightly detaching cancer cells with trypsin and scoring against CountBright (ThermoFisher) absolute counting beads on a flow cytometer.

#### CITE-Seq analysis of human tumors

For CITE-Seq, post tumor digestion, cells were incubated with Human TruStain FcX Receptor Blocking Solution to prevent non-specific antibody binding before staining with Zombie Aqua Fixable Viability Dye and anti-human CD45 antibody in PBS/2%FCS/2mM EDTA/0.01% sodium <u>azide</u> and incubated for 25 min on ice in the dark. Live CD45^+^ and CD45-cells were sorted on a BD FACSAria Fusion. CD45^+^ and CD45^-^ cells were pelleted and resuspended at 1×10^∧^3 cells/ml in 0.04%BSA/PBS buffer before mixing in an 8:2 CD45+:CD45-ratio and loaded onto the Chromium Controller (10X Genomics) to generate 5′ v1.1 gel beads-in-emulsions (GEM). Pooled 8:2 CD45+:CD45-cells were resuspended in Cell Staining Buffer (BioLegend) and stained with a pool of 137 TotalSeq-C antibodies (Table) according to the manufacturer’s protocol before loading onto the Chromium Controller (10X Genomics) for GEM generation. The cDNA libraries were generated using all or a subset of Chromium Next GEM Single Cell 5′ Library Kit for gene expression (GEX), Chromium Single Cell V(D)J Enrichment kit (10X Genomics) for T cell receptor (TCR), and Chromium Single Cell 5′ Feature Barcode Library kit for antibody derived tag (ADT) according to the manufacturer’s instructions. The libraries were subsequently sequenced on a Novaseq S4 sequencer (Illumina) to generate fastqs with the following mean reads per cell: 42,000 (GEX), 34,000 (TCR), and 5,700 (ADT). For multimodal clustering and analysis, CLR normalization followed by weighted nearest neighbor (WNN) clustering was performed using the Seurat package in R. Naïve and Exhaustion scores were generated using the percentile rank method as mentioned above, but with protein (ADT) markers-Naïve : CD62L, CD45RA, IL7RA; Exhaustion: PD-1, CTLA-4, CD38, CD39. Progenitor and Effector scores were generated from human orthologs of murine gene signatures (top 20 genes by log fold change) in Daniel et al. (*12*), removing those genes that have no expression in this dataset or no human equivalent. To compare cluster identities across datasets, we obtained gene signatures from the DEGs for each of our clusters and mapped their expression on to each major subset in a published pan-cancer intratumoral T cell atlas(*62*).

Q2 and Q4 gene signatures were generated by DEG of all intratumoral CD8 T cells belonging in these groups with a log fold change threshold of 0.25 and filtered by a p-value <0.01. For analysis of expression of various markers (*JUN, IFNG,* Exhaustion score) between Q2 and Q4, we select for those samples with at least 50 cells in a group to offer reliable metrics. This filters out certain patient samples when quantifying the same metrics within Tex^Early^ and Tex^Late^ subsets.

#### Live 2-photon imaging of tumor slices and image analysis

Live imaging of tumor slices was performed on a custom-made 2-photon microscope as previously described(*1*). Briefly, 1 million CD2dsRed; OT-I; Cd69-TFP or control CD2dsRed; OT-I CD8 T cells were retro-orbitally injected into WT mice bearing MC38chOVA tumors injected 5-7d earlier and harvested 7-10d after T cell injection. Slices for imaging were generated as described above for the ex vivo slice culture assay. Slices were placed in a custom-made perfusion chamber and imaged under oxygenated and temperature-controlled perfusion of RPMI 1640, as described previously(*1*). Dual laser excitations at 825nm and 920nm were used to excite the requisite fluorophores. Image analysis was performed on Imaris (BitPlane) with custom-made plugins developed on Matlab (Mathworks) and Fiji. Surfaces were generated on CD8 T cells and in both CD2dsRed; OT-I and CD2dsRed; OT-I; Cd69-TFP bearing slices and the corresponding levels of the former in the 515-545nm range PMT were used to gate on TFP^hi^ vs. TFP^lo^ OT-Is.

Cell tracking was performed on Imaris and corresponding cell positions imported to Matlab for further analysis to fit the persistent random walk model (PRWM) to the cell trajectories(*75*) using the method of overlapping intervals (*76*). Briefly, the mean squared displacement (MSD) for a cell for given time interval *t_i_* was obtained from the average of all squared displacements *x_ik_* such that

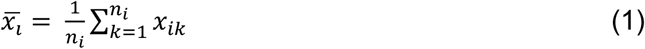

and

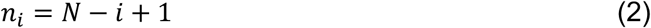

where *n_i_* is the number of overlapping time intervals of duration *t_i_* and N the total number of time intervals for the experiment. Mathematically, the persistent random walk model can be written as

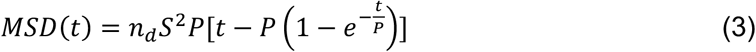

where S is the migration speed and P is the persistence time. The motility coefficient is given as

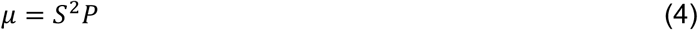

where *n_d_* is the dimensionality of the random walk (in this case *n_d_* =3). We fitted the PRWM in 3D to obtain estimates of speed, persistence time and motility of each cell track by non-linear regression.

#### Statistical Analysis

Statistical analysis was done in GraphPad Prism or in R. For testing null hypothesis between two groups, either Student’s t tests and or the non-parametric Mann-Whitney U tests were used, depending on the number and distribution of data points. Likewise, for testing null hypotheses among 3 or more groups, ANOVA or non-parametric tests were performed, followed by post-hoc Holm-Sidak’s test, correcting for multiple comparisons. Unless otherwise mentioned, data are representative of at least 2 independent experiments.

### Supplementary Figures

**Fig. S1:**
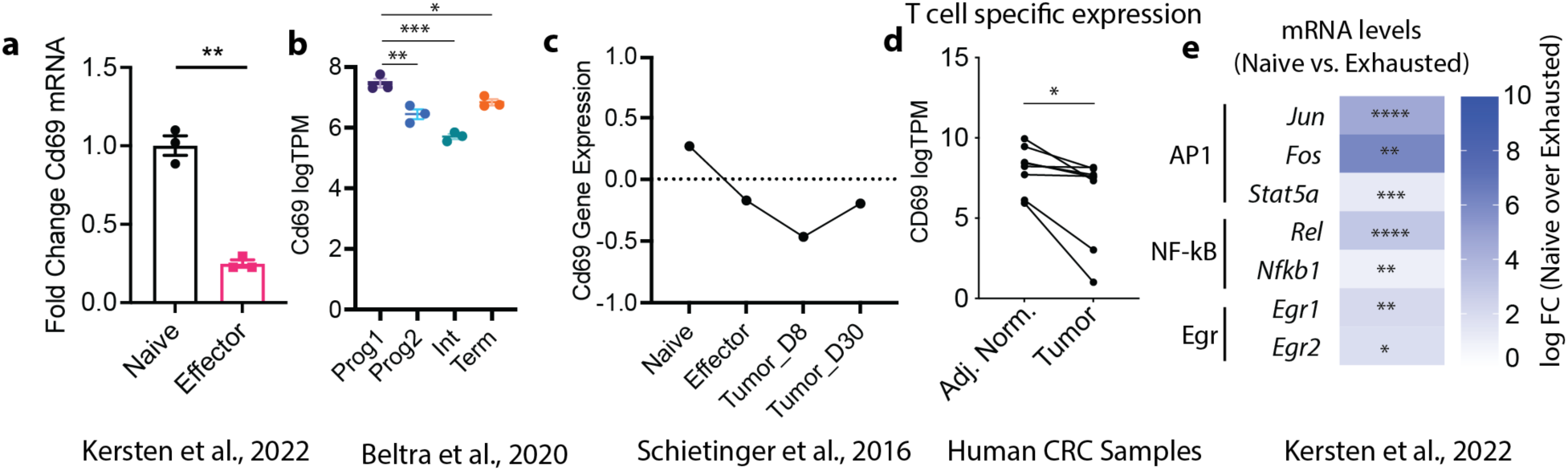
Resting Cd69 mRNA decreases with T cell differentiation towards exhaustion. Cd69 mRNA expression in **(a)** Naïve vs. in vitro generated (stimulation with Dynabeads followed by rest in IL-2 containing media) effector CD8 T cells from published RNASeq data(*23*); **(b)** Progenitor and exhausted T cell subsets from published data (*11*), **(c)** among Naïve, Effector, D8 tumor and D30 tumor infiltrating T cells from other published data (*8*), **(d)** in conventional T cells sorted from tumor and adjacent normal regions of human colorectal cancer patients; **(e)** depressed mRNA expression of factors associated with the Cd69 transcription in Naïve vs. Exhausted CD8 T cells from previous work (*23*) (symbols indicate FDR adjusted p-values). Plots show mean +/− SEM (a, b) p-values obtained by unpaired (a) and paired Student’s t test (d), and by ANOVA followed by post-hoc Tukey test. *p< 0.05, **p<0.01, ***p<0.001, ****p<0.0001.

**Fig. S2:**
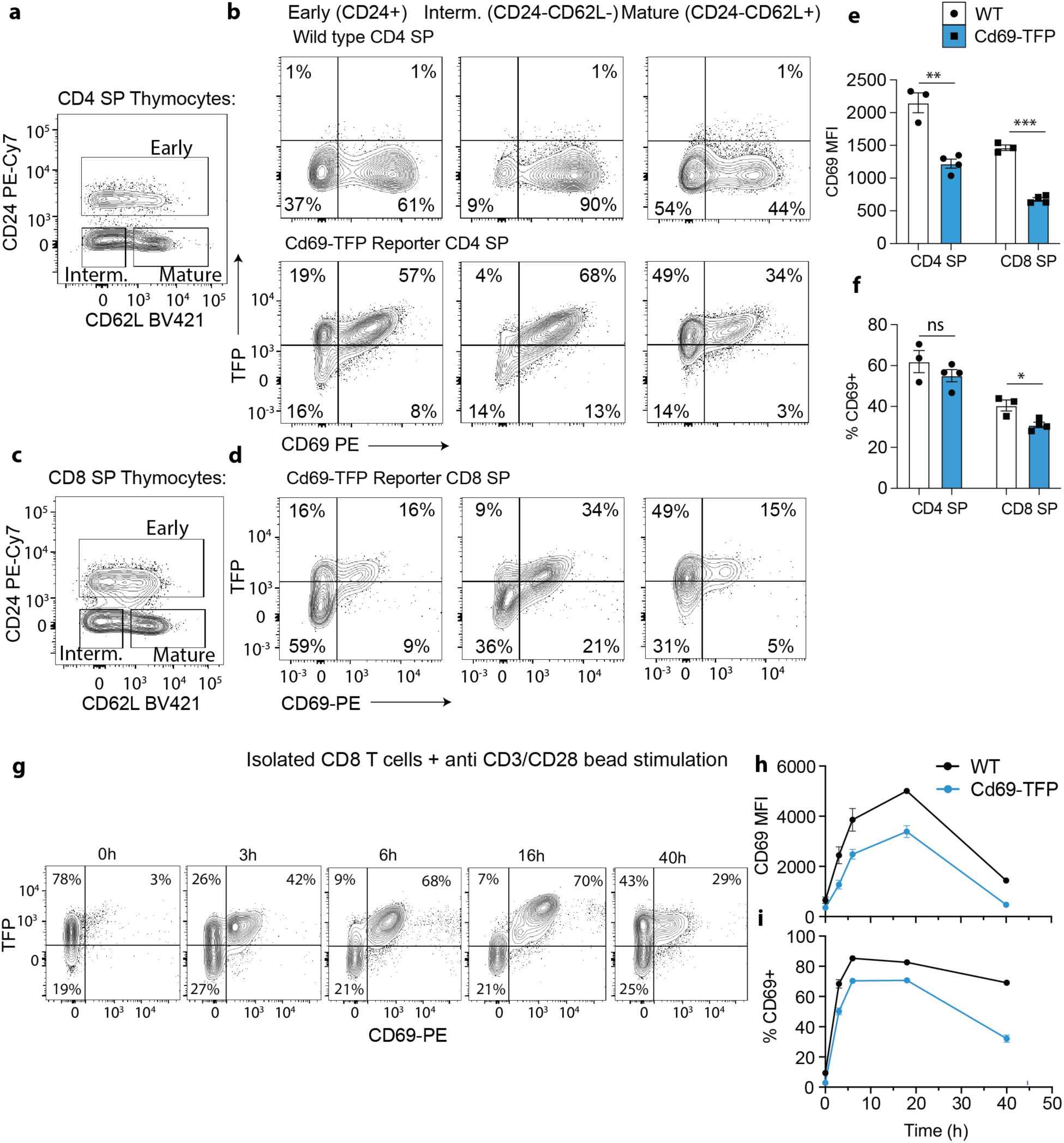
Cd69 transcriptional reporter TFP is expressed along with surface CD69 protein in known contexts of TCR stimulation. **(a, c)** Representative flow cytometry plot of CD62L and CD24 expression in **(a)** CD4+CD8-(CD4 single positive or SP) and **(c)** CD8+CD4- or CD8 SP thymocytes to demarcate early, intermediate (interm.) and mature subsets; **(b, d)** corresponding flow cytometry plots of TFP and CD69 expression in these subsets with varying degree of maturity in **(b)** CD4 SP cells in Cd69-TFP mice along with WT mice as controls and **(d)** in CD8 SP cells in Cd69-TFP reporter mice (WT: circular points, white bar; Reporter: square points, blue bars); quantification of **(e)** CD69 MFI and **(f)** %CD69+ in CD4 SP and CD8 SP cells from WT and Cd69-TFP reporter mice; **(g)** Representative flow cytometry plots of TFP vs. CD69 of isolated CD8 T cells from lymph nodes and spleen? Of an unchallenged reporter mouse at different time points post stimulation with αCD3+αCD28 Dynabeads and corresponding quantification of **(h)** CD69 MFI and **(i)** %CD69+ in these stimulated CD8 T cells from WT and Cd69-TFP reporter mice. *p< 0.05, **p<0.01, ***p<0.001, ns = no significance.

**Fig. S3:**
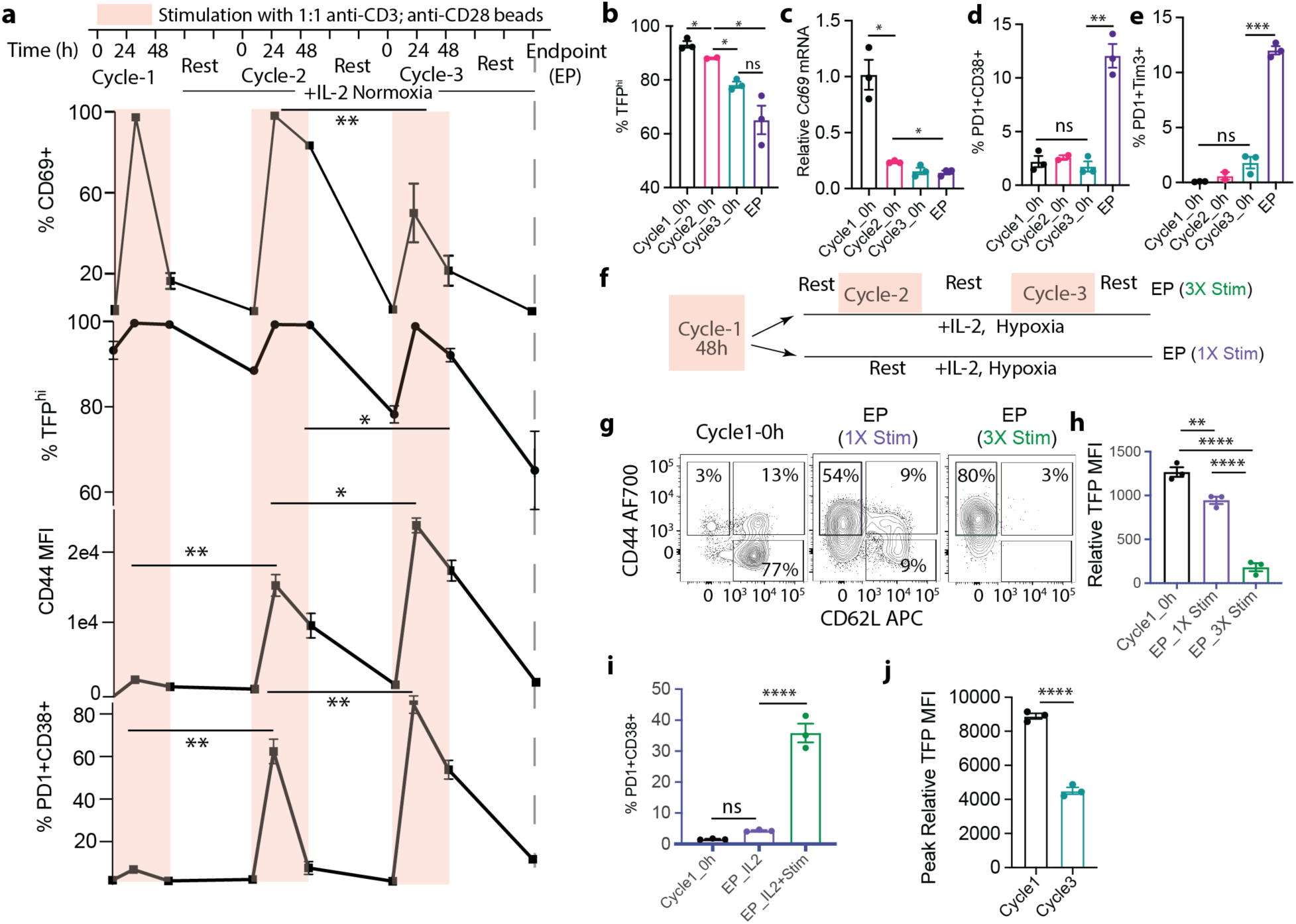
Repeated TCR stimulation drives down TFP with acquisition of exhaustion markers. **(a)** %CD69+, %TFP^hi^, CD44 MFI, %PD1^+^CD38^+^ of freshly isolated CD8 T cells through successive cycles of 48h stimulation and 72h resting in ambient oxygen (normoxia) + IL-2; **(b)** %TFP^hi^, **(c)** *Cd69* mRNA by qPCR and **(d)** %PD1^+^CD38^+^, **(e)** %PD1^+^Tim-3^+^ at the beginning of cycles 1, 2, 3 and endpoint (EP); **(f)** experimental schematic showing 1X Stim vs. 3X Stim conditions to parse the role of stimulation vs. IL-2 alone; **(g)** flow cytometry plots showing representative CD44 vs. CD62L profiles of CD8 T cells at the timepoints and conditions indicated; for the same experiment, **(h)** TFP (relative to WT control), **(i)** %PD1^+^CD38^+^ of CD8 T cells at the starting point (Cycle1_0h) and at endpoint (EP) either with 1X Stim followed by prolonged rest or 3X stim; **(j)** Peak Relative TFP and CD69 MFI between Cycle 1 and Cycle 3 in normoxia; Bar graphs represent mean +/− SEM; null hypothesis testing by ANOVA followed by post-hoc Holm-Šídák test; data representative of 2 independent experiments, each with 3 mice and technical duplicates/biological replicate at every assay point. *p< 0.05, **p<0.01, ***p<0.001, ****p<0.0001, ns = no significance.

**Fig. S4:**
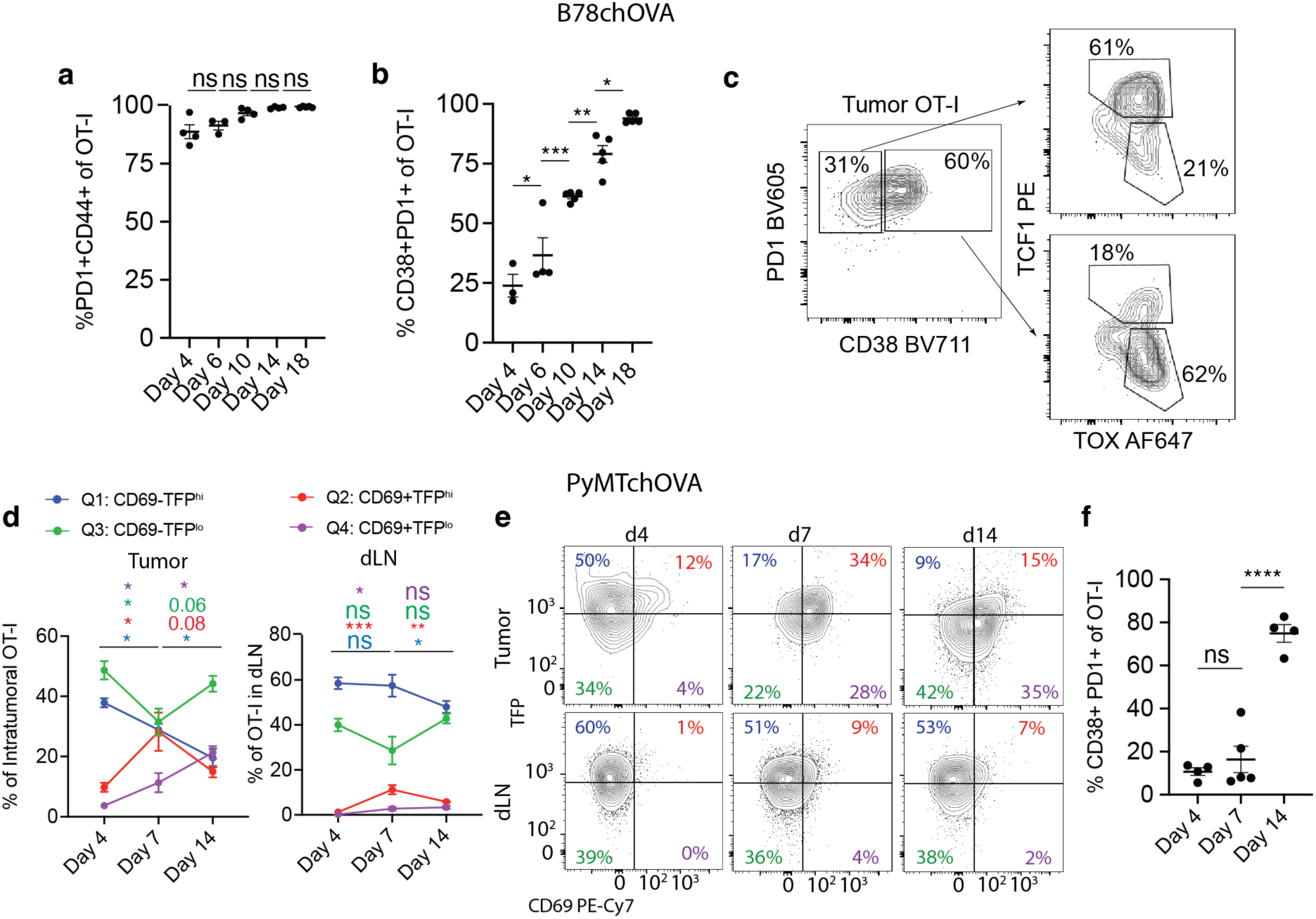
TFP decrease accompanies antigen-specific CD8 T cell residence time in TME: **(a)** %PD1+CD44+ and **(b)** %PD1+CD38+ of intratumoral OT-I CD8 T cells at different time points after adoptive transfer into WT mice bearing B78chOVA tumors; **(c)** Flow cytometry plots showing TCF1 and TOX expression in PD1+CD38- and PD1+CD38+ intratumoral OT-Is d14 post adoptive transfer in the same tumors; (**d)** CD69:TFP quadrant (Q1-Q4) distribution for OT-I T cells in the tumor and tdLN over time after adoptive transfer into PyMTchOVA mice bearing mammary tumors**; (e)** corresponding corresponding flow cytometry plots showing TFP and CD69 expression of the same adoptively transferred OT-I T cells; **(f)** %PD1+CD38+ of intratumoral OT-I CD8 T cells at different time points after adoptive transfer into PyMTchOVA mice bearing mammary tumors. Bar graphs show mean +/− SEM. *p<0.05, **p<0.01, ***p<0.001, ****p<0.0001, 0.05<p<0.1 are recorded, ns: p>=0.1, by one-way ANOVA with post-hoc Holm-Sidak multiple comparison test for a, b, f and 2-way ANOVA with post-hoc Tukey test between pair-wise timepoints in (d).

**Fig. S5:**
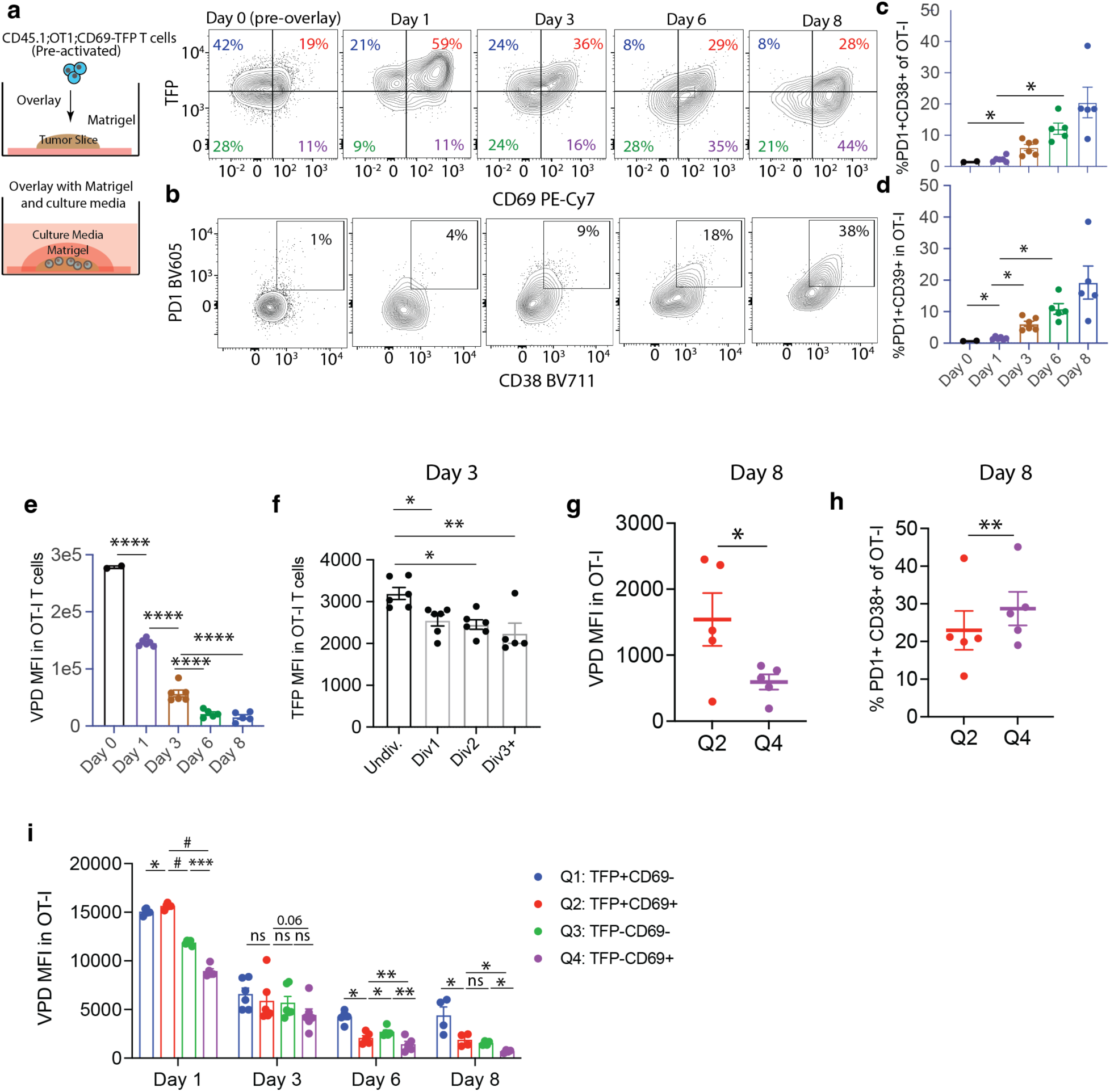
Ex vivo tumor slice overlay culture mimics OT-I differentiation in vivo: Schematic of tumor slice culture assay with representative flow cytometry plots of **(a)** TFP vs. CD69 and **(b)** PD1 vs. CD38 expression in slice-internal OT-I T cells from Day1-Day8, compared to Day0 (pre-overlay); **(c)** %PD1+CD38+, **(d)** %PD1+CD39+ and **(e)** Violet proliferation dye (VPD) MFI of Day 0 pre-overlay and slice-internal OT-I T cells at different time points after slice overlay (Day1-Day8); **(f)** bar graph showing TFP MFI of slice OT-I T cells by estimated number of divisions (>=3, 2, 1, or divided) at Day 3; **(g)** VPD MFI and **(h)** %PD1+CD38+ of slice-internal OT-Is from Q2 and Q4 at Day8; **(i)** VPD MFI by CD69:TFP quadrants over time in OT-I T cells recovered from within slices; Bar graphs show mean +/− SEM, null hypothesis testing by ANOVA and post hoc Holm-Šídák test (c-f), or paired t test in g, h, 2-way ANOVA and multiple comparison tests controlling for False Discovery Rates (i); data are representative of 2 independent experiments, each 5-6 slices/time point for each slice experiment and Day 0 pre-overlay samples in duplicate; TFP gated on OT-I control (without Cd69-TFP) CD8 T cells. *p< 0.05, **p<0.01, ***p<0.001, **** or #p<0.0001.

**Fig. S6:**
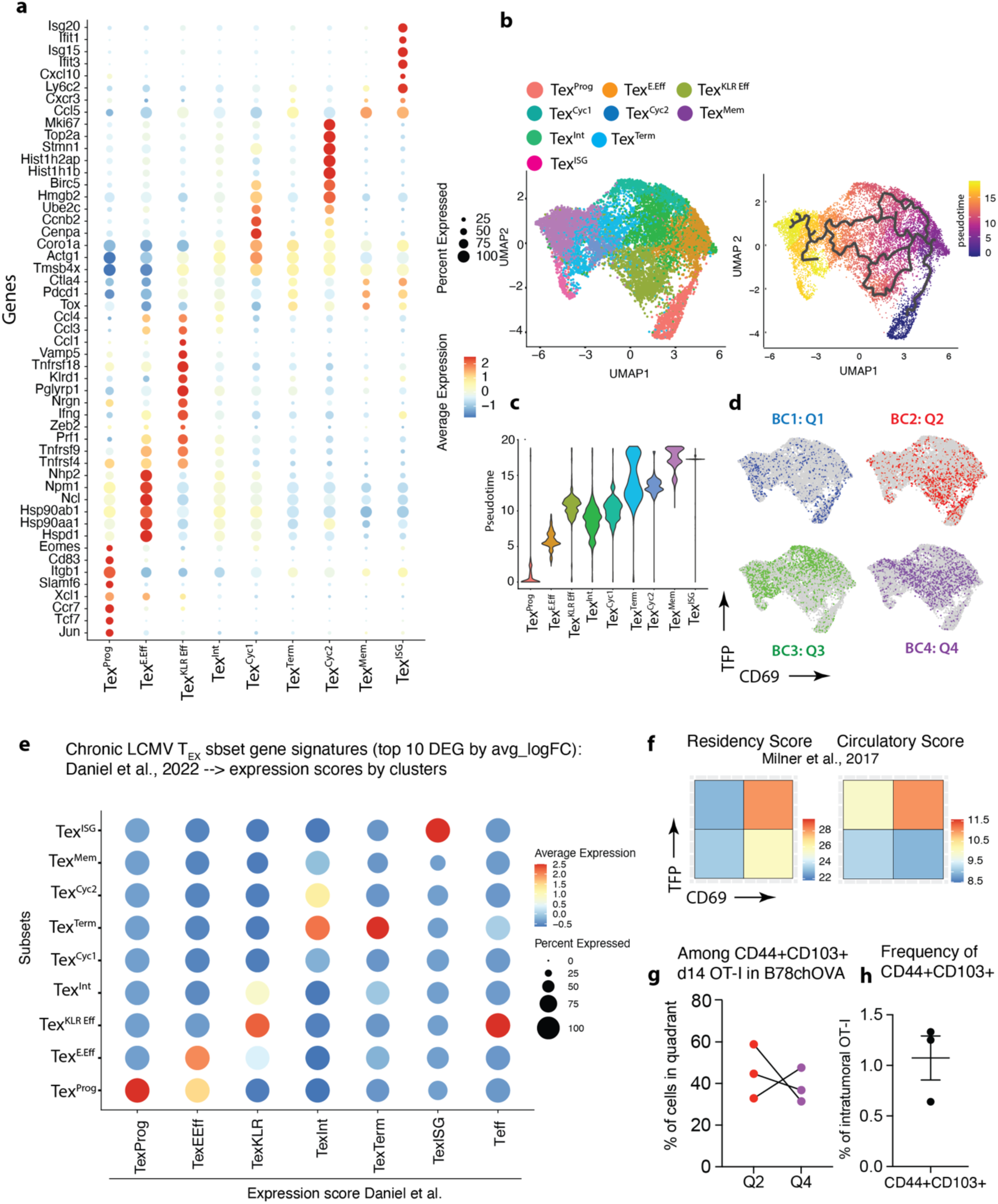
Gene expression based clustering of intratumoral OT-Is and relationship to Cd69 transcription. **(a)** Dotplot representation of differentially expressed and canonical T_EX_-associated genes across the computationally derived cell clusters from the scSeq of intratumoral Cd69-TFP:OT-I CD8 T cells at d12 post injection into B78chOVA tumor-bearing mice; **(b)** UMAP representation of the scSeq data color-coded by pseudotime derived from Monocle3 trajectory analysis and **(c)** Pseudotime spread of each cluster from the same analysis; **(d)** overlay of cells with barcodes corresponding to the sorted CD69:TFP clusters in the UMAP space; **(e)** Average expression of the signature scores from top 10 DEGs of LCMV-derived T_EX_ clusters from Daniel et al.(*12*) in the subsets obtained from this study; **(f)** Expression score derived from transcriptomic signatures of CD8 T cell residency vs. circulation, as defined by Milner et al., 2017; **(g)** distribution in Q2 vs. Q4 quadrants and **(h)** frequency of CD44+CD103+ T_RM_-like cells in adoptively transferred d14 OT-I T cells in B78chOVA tumors.

**Fig. S7:**
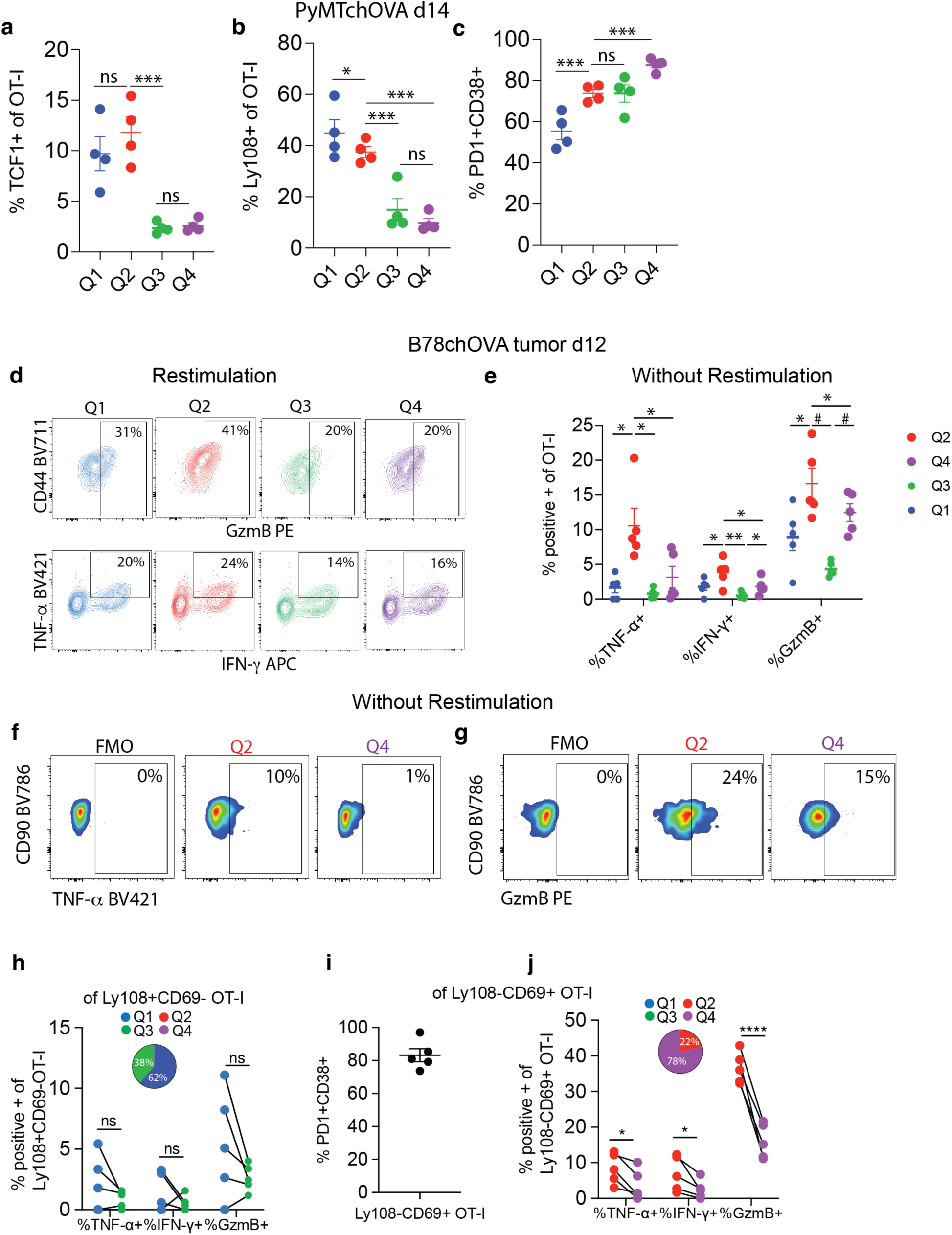
**(a)** % TCF1+, **(b)** %Ly108+, **(c)** %PD1^+^CD38^+^ of intratumoral OT-I T cells grouped by CD69:TFP quadrants in PyMTchOVA mammary tumors 14d post-adoptive transfer; **(d)** representative flow cytometry plots showing TNF-α, IFN-γ and GzmB expression in intratumoral OT-I T cells in B78chOVA tumors 12d post adoptive transfer by Cd69:TFP quadrants, assayed directly by intracellular staining without sort and restimulation; **(e)** quantification of TNF-α, IFN-γ and GzmB expression of the same intratumoral OT-I T cells, sorted by Cd69:TFP quadrants and restimulated for 3h with PMA/ionomycin; flow cytometry plots showing intracellular **(f)** TNF-α and **(g)** GzmB expression in intratumoral OT-I T cells 12d post adoptive transfer into B78chOVA tumors; cytokine and GzmB expression of Q2 vs. Q4 OT-I T cells within **(h)** Ly108+CD69-(*11*) and **(j)** Ly108-CD69+(*11*) subsets with inset pie chart showing the percentage of cells in each quadrant within the corresponding subsets; **(i)** %PD1^+^CD38^+^ among Ly108-CD69+ intratumoral OT-Is; graphs represent mean +/− SEM; null hypothesis testing by RM ANOVA followed by post-hoc paired t-tests corrected for multiple comparisons; *p< 0.05, **p<0.01, ***p<0.001, ****p<0.0001, ns = no significance, #p<0.001 in (e).

**Fig. S8:**
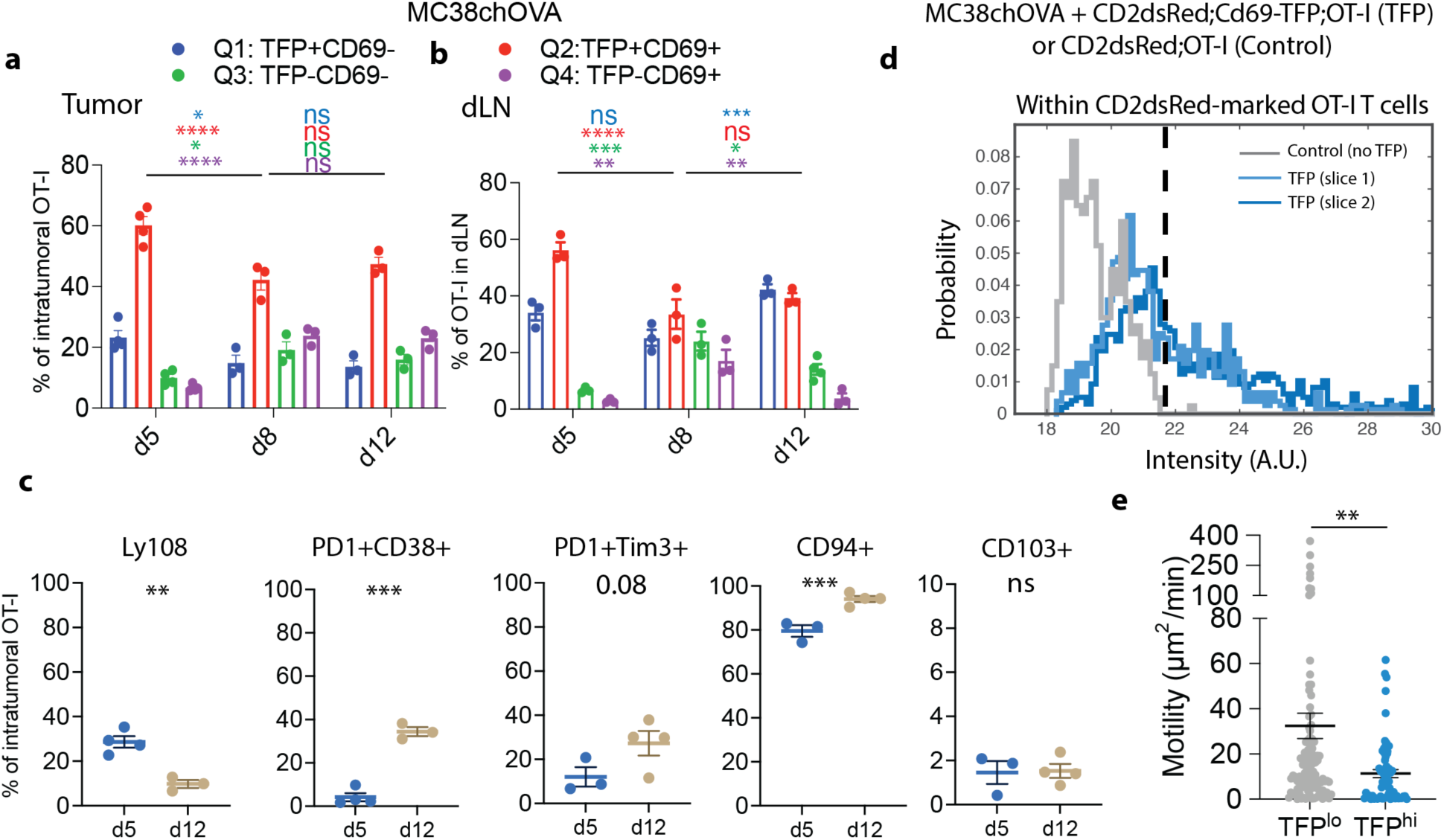
Bar graphs showing TFP:CD69 quadrant distribution among OT-I CD8 T cells in **(a)** tumors and **(b)** corresponding tdLN at d5, d8, d12 post T cell injection into MC38chOVA tumor-bearing mice (n=3-4 mice per group respectively); **(c)** expression of surface markers of progenitor, exhausted, effector and T_RM_-like states in intratumoral OT-Is (d5 and d12) in MC38chOVA tumors; **(d)** Representative histograms of channel intensity within OT-I T cells in live tumor slices to find TFP^hi^ cells using CD2dsRed and CD2dsRed;Cd69-TFP OT-Is; **(e)** overall 3D motility of TFP^hi^ vs. TFP^lo^ intratumoral OT-Is d8 post adoptive transfer within live MC38chOVA tumor slices; bar graphs show mean +/− SEM, null hypothesis testing by Mann-Whitney U test (e), 2-way ANOVA with post-hoc Tukey test between pair-wise timepoints (a, b) and unpaired t tests (c). *p< 0.05, **p<0.01, ***p<0.001, ****p<0.0001, ns = no significance.

**Fig. S9:**
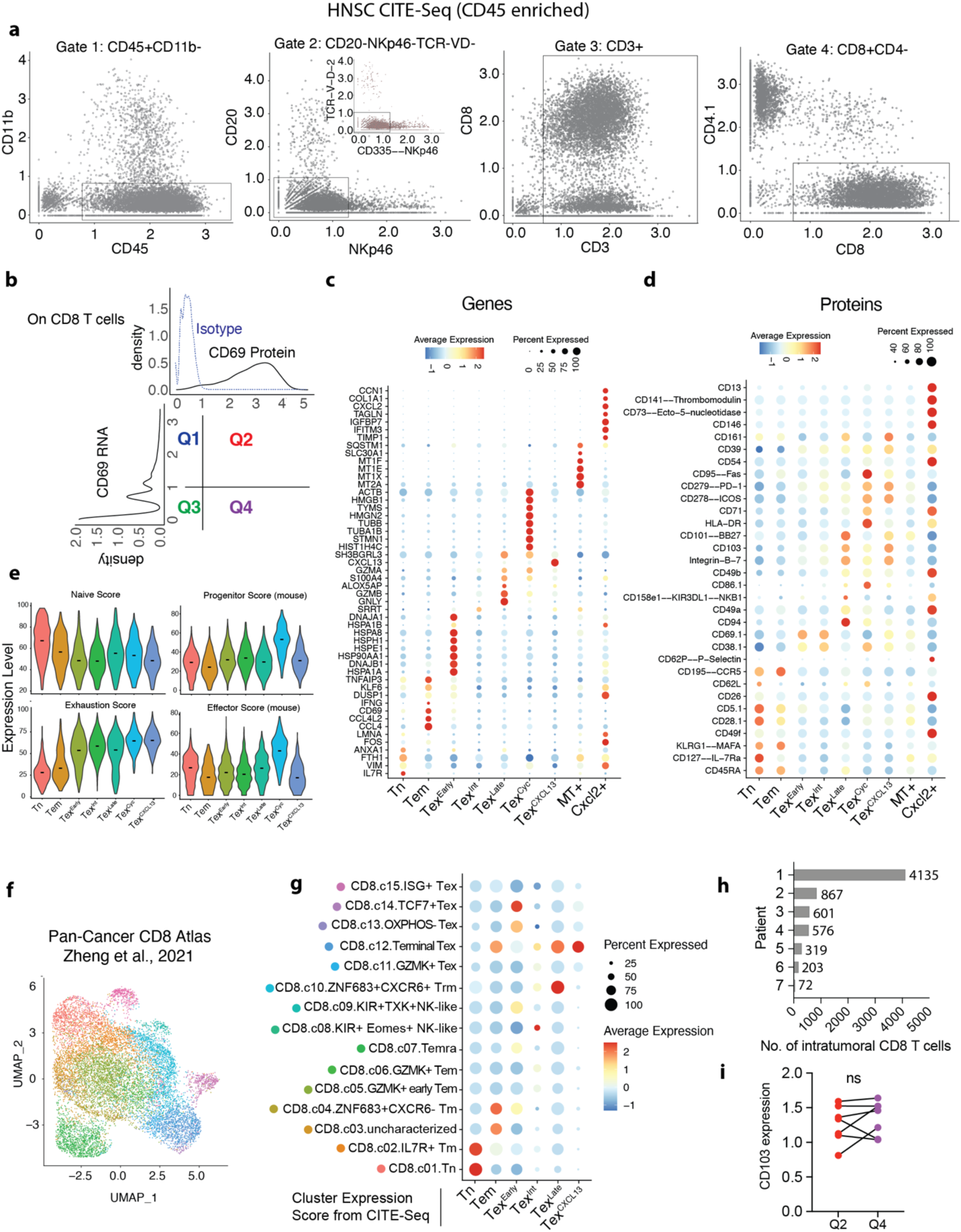
CITE-Seq of HNSC tumor allows mapping of Cd69 RNA:CD69 Protein quadrants onto CD8 T cell phenotypes. **(a)** Gating scheme of CD45-enriched CITE-Seq from HNSC patient data using protein markers to isolate a pure CD8 T cell population; **(b)** Gating of the CD8 T cell population into CD69 Protein: CD69 RNA quadrants (Q1-Q4); **(c)** DEGs and **(d)** DE antibody-tagged surface proteins for the computationally derived subsets obtained through multimodal clustering using both protein and RNA; **(e)** progenitor and effector scores derived from (*12*) along with naïve and exhaustion scores (based on antibody-derived tags as in Fig. 6c). **(f)** UMAP representation of computationally-derived subsets among CD8 T cells in a pan-cancer T cell atlas(*62*), color-coded subset labels are as shown in (g); **(g)** DEGs from each subset in the CITE-Seq data (avg_logFC>0.5, adj_pvalue <0.01) were used to generate gene signature scores and mapped onto the major pan-cancer CD8 T cell subsets; **(h)** Number of CD8 T cells recovered by gating in a from each patient in the CITE-Seq data; **(i)** CD103 (antibody-tagged) expression among Q2 and Q4 (gated as in b) intratumoral CD8 T cells from CITE-Seq, ns: no significance by paired t-test.

**Fig. S10:**
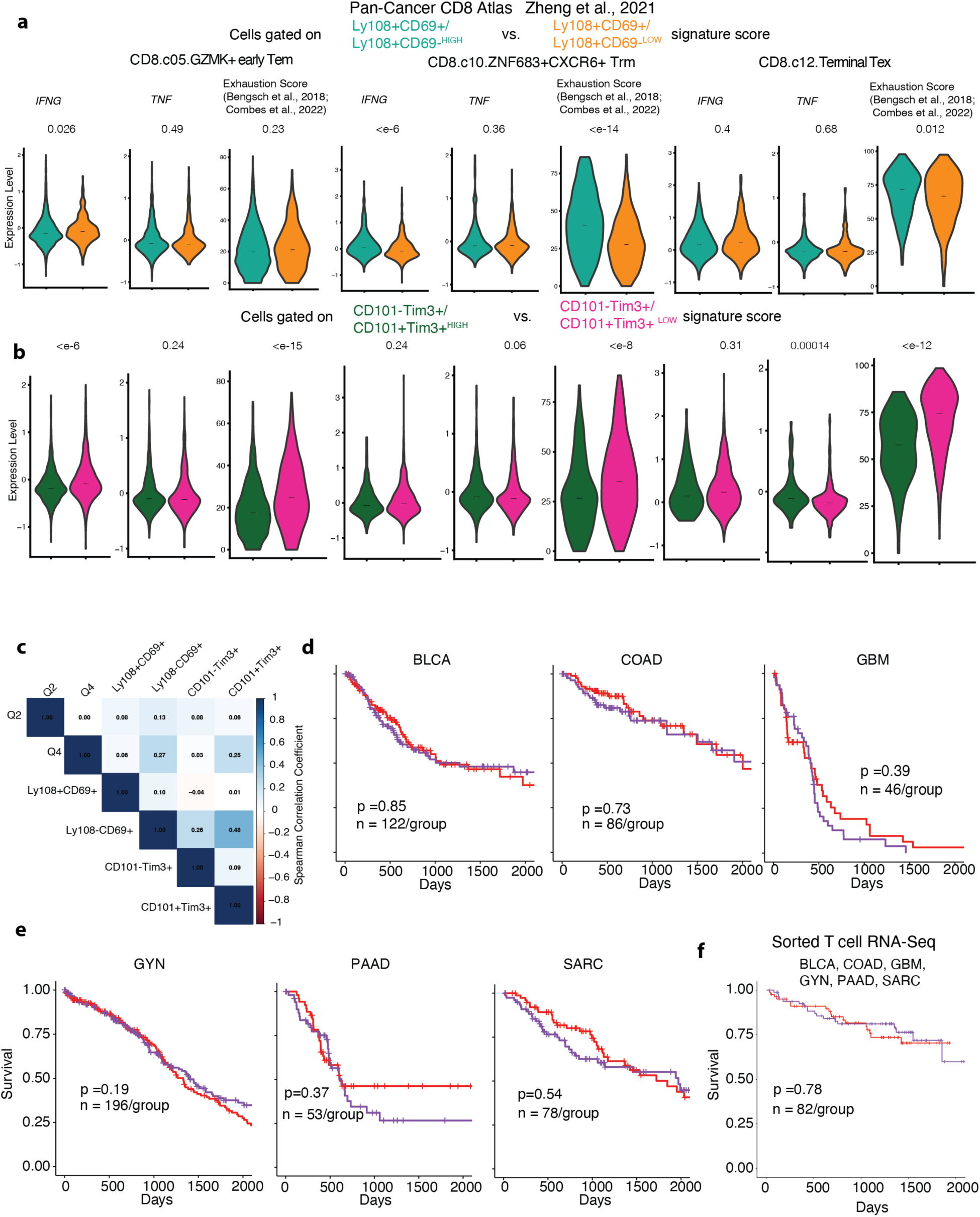
**(a, b)** expression of *IFNG*, *TNF* and Exhaustion signature score within the Early Tem, CXCR6+ Trm and Terminal Tex clusters from the Pan-Cancer CD8 atlas (Zheng et al., 2021), grouped by signatures corresponding to **(a)** the ratio of Ly108+CD69+ and Ly108-CD69+ expression scores (Signature 1 and Signature 7 from (*11*), and **(b)** the ratio of expression scores for the CD101-Tim3+ effector and CD101+Tim3+ terminal subsets (*13*); **(c)** Spearman correlation matrix showing the relationship among the used gene signatures, Q2 and Q4 from this study, Signature 1 (Ly108+CD69+), Signature 7(Ly108-CD69+) from (*11*), CD101-Tim3+ and CD101+Tim3+ from (*13*); Kaplan-Meier survival curves from a quality controlled subset of TCGA data (*63*) corresponding to **(d)** BLCA, COAD, GBM, **(e)** GYN, PAAD and SARC indications, patients stratified by the ratio of Q2 and Q4 expression scores; **(f)** Kaplan-Meier survival curves from a cohort of patients of combined BLCA, COAD, GBM, GYN, PAAD, SARC indications, with patients stratified by the ratio of Q2 and Q4 expression scores from bulk RNA-Seq of sorted T cells(*63*) number of patients per group (after stratification) and p-value for log-rank tests are noted in each panel.

## Notes

### Competing Interest Statement

The authors have declared no competing interest.

### Summary of Updates

Format has been changed from 5 figure to 7 figure. This version goes into more depth on the difference between Q2 and Q4 states delineated by Cd69 RNA (TFP) expression, without delving into specific surface marker identity of the most potent effectors. Title of the manuscript has been modified concordant with this change in focus. Fig. 1 and 2 are enhanced and modified versions of the previous Fig. 1; Fig. 3 is the previous Fig. 2 with significant enhancements; Fig. 4 and Fig. 5 are enhanced versions of previous Figs. 3 and 4; Fig. 6 is an altered version of previous Fig. 5; Fig. 7 contains new, additional human data. Supplementary Figs. have been accordingly altered. Graphical abstract has been added.

